# Environmental plasticity and colonisation history in the Atlantic salmon microbiome: a translocation experiment

**DOI:** 10.1101/564104

**Authors:** Tamsyn M. Uren Webster, Deiene Rodriguez-Barreto, Giovanni Castaldo, Peter Gough, Sofia Consuegra, Carlos Garcia de Leaniz

## Abstract

Microbial communities associated with the gut and the skin are strongly influenced by environmental factors, and can rapidly adapt to change. Historical processes may also affect the microbiome. In particular, variation in microbial colonisation in early life has the potential to induce lasting effects on microbial assemblages. However, little is known about the relative extent of microbiome plasticity or the importance of historical colonisation effects following environmental change, especially for non-mammalian species. To investigate this we performed a reciprocal translocation of Atlantic salmon between captive and semi-natural conditions. Wild and hatchery-reared fry were transferred to three common garden experimental environments for six weeks: standard hatchery conditions, hatchery conditions with an enriched diet, and simulated wild conditions. We characterised the faecal and skin microbiome of individual fish before and after the environmental translocation, using a BACI (before-after-control-impact) design. We found evidence of extensive plasticity in both gut and skin microbiota, with the greatest changes in alpha and beta diversity associated with the largest changes in environment and diet. Microbiome richness and diversity were entirely determined by environment, with no detectable historical effects of fish origin. Microbiome structure was also strongly influenced by current environmental conditions but, for the first time in fish, we also found evidence of colonisation history, including a number of OTUs characteristic of captive rearing. These results may have important implications for host adaptation to local selective pressures, and also highlight how conditions during early life can have a long-term influence on the microbiome and, potentially, host health.

## Introduction

Microbiome structure is determined by a complex series of delicately balanced interactions with the host, the environment and amongst microbiota (Suzuki 2017; Vellend 2016; Walter & Ley 2011). Unlike the host genome, the metagenome is very dynamic and readily influenced by host-specific and environmental changes (Chen et al. 2018; Davenport et al. 2017). Host-associated microbial communities are able to rapidly respond to local selective pressures due to their short generation times, rapid mutation rates, large population sizes and high levels of phenotypic plasticity and intra-community gene flow (Walter & Ley 2011). Given the critical and wide-ranging influence of the microbiome on host health and fitness (Davenport et al. 2017; Koskella et al. 2017), extensive metagenomic plasticity may also enhance host adaptive phenotypic plasticity (Alberdi et al. 2016; Suzuki 2017). This may play a vital role in driving host adaptation to environmental change, and may even contribute to population-level divergence and local adaptation (Alberdi et al. 2016; Walter & Ley 2011). For example, intestinal microbiota can enhance the digestion of novel food sources and the metabolism of dietary toxins, increase drought and thermal tolerance, and increase resistance to local pathogens (Alberdi et al. 2016; Chevalier et al. 2015; Macke et al. 2017).

While microbial communities often show extensive plasticity in response to environmental change, historical processes may potentially also affect the microbiome. The order and timing in which microorganisms arrive in the community can impact microbiome structure, even under otherwise identical environmental conditions (Sprockett et al. 2018; Vellend 2016). Existing colonisers may restrict and/or modify the ecological niches available to subsequent arrivals, influencing their establishment success (Fukami 2015). For example, environmental stress or antibiotic treatment which disturbs the microbiome may allow resistant taxa to flourish in the absence of wider competition, delaying the restoration of the original complex community (Foster et al. 2017; Yassour et al. 2016). During the early stages of microbiome colonisation and establishment, microbial composition is particularly dynamic and readily influenced by variation in the surrounding environment and microbial seeding communities (Korpela et al. 2018; Stephens et al. 2016). Therefore historical effects on microbiome structure due to environmental variation during early life may be particularly important (Gensollen et al. 2016b; Sprockett et al. 2018). Early life stress can have a lasting detrimental effect on the microbiome and health outcomes in mammalian hosts (Foster et al. 2017), but the possibility of conditioning the microbiome during early life to improve lasting host health and disease resistance could also have huge therapeutic benefit (Borre et al. 2014). However, overall, little is known about the extent to which historical effects may shape the microbiome, especially for non-mammalian species.

Atlantic salmon (*Salmo salar*) is one of the most commercially important fish species and is also a keystone species for freshwater habitats (Griffiths et al. 2011). Populations of Atlantic salmon display local adaptation, and are often threatened in their natural range (Garcia de Leaniz et al. 2007). The Atlantic salmon gut and skin microbiome is known to be strongly influenced by environmental conditions, especially diet, salinity and season, as well as developmental stage (Dehler et al. 2017; Gajardo et al. 2016a; Llewellyn et al. 2016; Lokesh & Kiron 2016; Zarkasi et al. 2014), and there is also substantial microbiome variation between salmon populations, especially between wild and hatchery-reared fish, reflecting diet and environmental differences (Uren Webster et al. 2018). This suggests there may be considerable scope for microbiome plasticity in response to environmental variation, but may also indicate the potential for lasting historical effects reflecting colonisation history and early life experience, and makes Atlantic salmon an ideal species in which to examine these factors. Understanding the scope for plasticity and historical effects is essential for the management of this species both in aquaculture and the wild. For example, microbiome plasticity and/or historical effects may influence acclimation to environmental challenges, act as a driver of local adaptation, or provide a potential mechanism for improving disease resistance in aquaculture. However, little is known about the scope for plasticity following environmental change, and/or whether historical colonisation effects may continue to influence microbiome diversity and structure. Therefore, to investigate this, we used a reciprocal translocation experiment of salmon fry between hatchery and natural conditions and a BACI (before-after-control-impact) experimental design to test for gut and skin microbiome plasticity related with development and environment/diet, as well as potential lasting historical signatures of origin.

## Materials and Methods

### Reciprocal translocation experiment

Sixty wild Atlantic salmon fry (∼6 months post hatch) were captured using electrofishing from the Aber Bran, a tributary of the river Usk (Wales; lat: 51.954, long: −3.477) and transported a short distance (∼8 km) to the NRW Cynrig Fish Culture Unit (Brecon, Wales). Sixty Atlantic salmon fry (6 months post hatch) were also obtained from the hatchery; these fish originated from a 3:3 male:female cross between wild-caught parents from the river Taff (Wales) that had been maintained under hatchery conditions. Hatchery fish were adipose fin clipped to differentiate them from wild fish, a procedure that does not cause adverse effects (Roberts et al. 2014). All wild and hatchery fish were measured (fork length) and photographed using a Canon DS126151 400D EOS digital camera, using a 18-55 mm lens. A sample of skin-associated mucus was collected by swabbing the left side of each fish back and forth along the entire length of the lateral line five times using Epicentre Catch-All(tm) Sample Collection Swabs (Cambio, Cambridge, UK), and gut samples were collected by gently pressing the abdomen of each fish and swabbing the expelled faeces. Duplicate 50 ml water samples were also collected from the river and hatchery. All samples were stored at −80°C prior to DNA extraction.

Twenty wild and 20 hatchery individuals were randomly assigned to each of the three experimental environments (hatchery, enriched diet, natural) using a common garden design (Figure 1). The first experimental group was maintained in standard hatchery conditions. Fish were housed in a 500 L black plastic tank supplied with aerated, flow-through, filtered river water (∼10 L/min) and fed a standard commercial Salmonid feed (Skretting) at a rate of 3% body weight/day. The second group were housed in identical conditions to the first group, but were fed the standard hatchery diet enriched with daily addition of 5 g of an invertebrate mix (33% natural invertebrates harvested from the Afon Cynrig, 33% bloodworm and 33% daphnia (both JMC Aquatics, Sheffield, UK)). The third experimental group consisted of near natural conditions, whereby salmon were transferred to a 30m leat of the Afon Cynrig, another tributary of the river Usk (lat: 51.928, long: −3.358). The leat was isolated by upstream and downstream fry screens to prevent fish movement, contained natural substrate, was fed by natural river water, and received no dietary supplementation. The experimental treatments lasted for six weeks, after which fish were recaptured, and sampled for a second time using an identical procedure to the one described for the first sampling point (photograph, fork length, weight, skin swab, faecal swab). Duplicate water samples were collected from the hatchery tank, enriched tank and leat. Water temperature in each of the experimental tanks and the leat ranged from 13.5-16.5 °C during the experiment reflecting ambient conditions.

**Figure 1.**
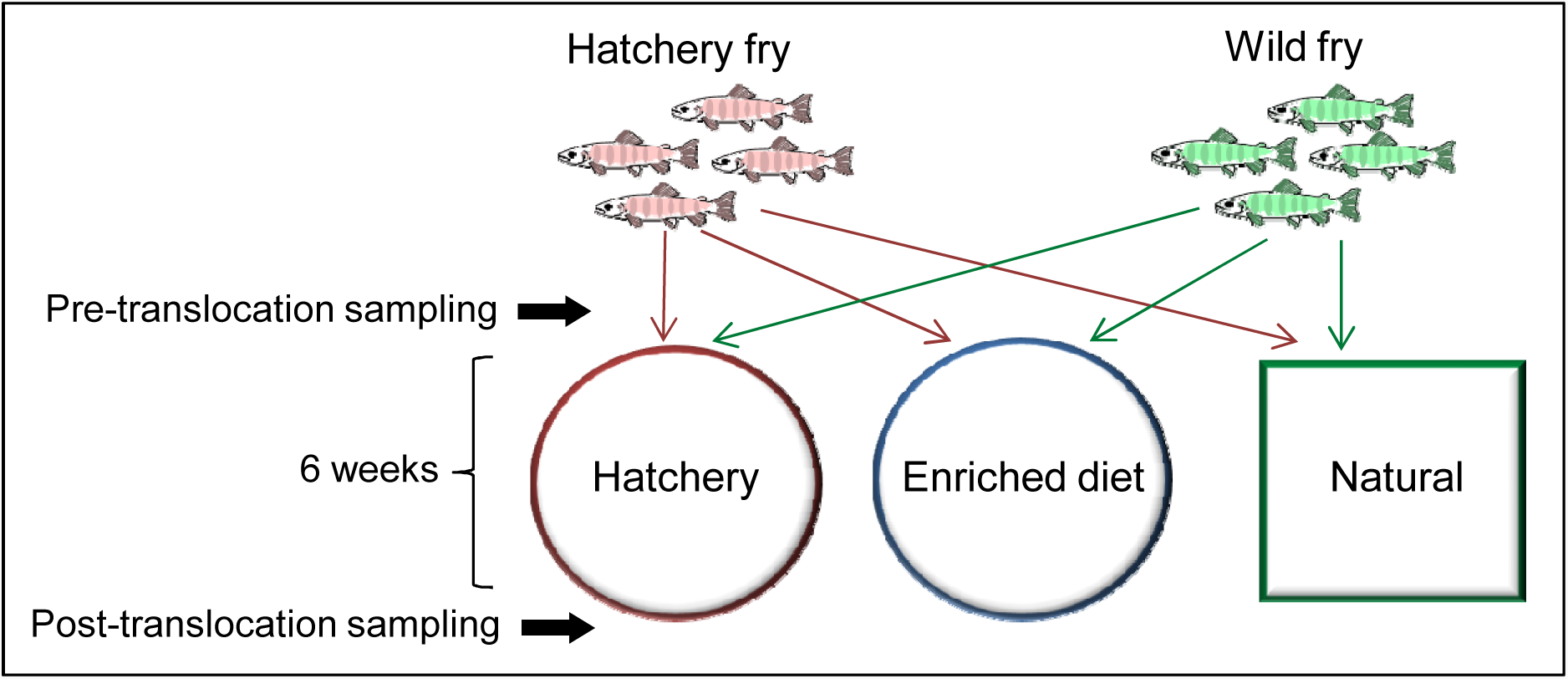
Before-after-control-impact (BACI) design. Hatchery- and wild-origin salmon fry were translocated to three experimental environments, employing a common garden design. Individual fish were matched at the pre- and post-translocation sampling based on unique pigmentation marks.

Photographs were used to visually match individual fish at the first and second sampling points, allowing us to conduct a matched before-and-after microbiome analysis. Each fish was identified based on the number, shape and spacing of its parr marks, as juvenile salmonids can be identified based on a unique pattern of pigmentation (Donnelly & Whoriskey 1993; Garcia de Leaniz et al. 1994). All matches were independently corroborated by two researchers, and eight individuals from each of the six experimental groups (wild to hatchery; hatchery to hatchery; wild to enriched; hatchery to enriched; wild to natural; hatchery to natural) were positively identified at both time-points and used for further analyses. Specific growth rate ((Ln(length_2_)-Ln(length_1_))/time x100; (Hopkins 1992)) and Fulton’s condition factor ((weight/length^3^) × 100; (Froese 2006) were calculated for each individual fish.

### 16S rRNA amplicon sequencing and bioinformatics

Microbiome analysis was performed for the skin and gut samples at the two time-points for 48 individual fish, as well as five water samples in duplicate. DNA was extracted from the skin mucus and faecal samples using the MoBio PowerSoil® DNA Isolation Kit (Qiagen) according to the manufacturer’s instructions, with an additional incubation step of 10 minutes at 65°C prior to bead beating. Water samples were centrifuged at 5000xg for 1 hour at 4°C and the DNA was extracted from the pellet using the same method. 16S library preparation was performed as described previously (Uren Webster et al. 2018) amplifying the V4 hypervariable region of the bacterial 16S gene using the primers 341F and 785R, and libraries were sequenced across two lanes of an Illumina MiSeq.

Quality filtering of raw sequence reads (removal of adaptor sequences and poor quality bases from the 3’ end (Q<20)) was performed using Trimmomatic (Bolger et al. 2014), before analysis using mothur v1.39 (Kozich et al. 2013) and Qiime v1.9 (Caporaso et al. 2010). Forward and reverse reads were concatenated, filtered to retain amplicons of the target size range, then aligned to the Silva seed reference database (version 128)(Quast et al. 2013). Further noise reduction of contigs with high quality alignments was performed using mothur’s pre-clustering algorithm, then potential chimeras were identified and removed using UCHIME (Edgar et al. 2011) within mothur. Taxonomic classification of contigs was performed using the Silva reference taxonomy (Quast et al. 2013) and mitochondrial, eukaryote and chloroplast sequences were removed. Contigs were clustered into operational taxonomic units (OTUs) based on 97% overall sequence similarity. Singleton OTUs were removed from the dataset, and all gut, skin and water samples were subsampled to an equal depth of 19,389 reads. Microbial alpha diversity analysis was performed based on Chao1 richness and Shannon diversity, while structural analysis (microbial beta diversity) was based on community distance matrices calculated using the Bray-Curtis dissimilarity index (Bray & Curtis 1957), which accounts for OTU presence/absence and relative abundance. Functional analysis of community structure was performed based on metagenomic predictions made using PICRUSt (Langille et al. 2013). The OTU table produced using mothur was converted to a compatible biom format based on Greengenes reference taxonomy v13_8 (DeSantis et al. 2006) using biom convert (McDonald et al. 2012). PICRUSt was then used to normalise OTUs, predict metagenomes based on KEGG orthologs and collapse functional predictions to KEGG pathway level. Further visualisation and statistical analysis based on KEGG pathways was then performed in STAMP v2.1.3 (Parks & Beiko 2010).

### Statistical analysis

All statistical analyses were performed using R (v3.4.3; (R_Core_Team 2014)). To investigate differences in the specific growth rate (SGR) of individually identified fish during the course of the translocation experiment, and in final condition factor (K), linear models including the fixed factors environment (hatchery, enriched, natural), origin (hatchery, wild), and their interaction, were constructed. The best fitting models were selected based on Akaike Information Criterion (AIC) values using the *step* function (Table S1).

To identify initial differences in alpha diversity (Chao1 richness and Shannon diversity) at the pre-translocation sampling point, linear models were constructed including origin (hatchery/wild) and fish size (length) as fixed factors. The effects of experimental environment (hatchery/enriched/natural), origin, an environment:origin interaction and SGR on the specific change in alpha diversity for matched individuals during the course of the translocation experiment were also examined. Finally, linear models were constructed to investigate the effect of environment, origin, an environment:origin interaction and size (length), as well as the pre-sampling alpha diversity value for matched individual fish, on final (post-translocation) alpha diversity values (Chao1 richness and Shannon diversity). In each case reduced linear models were selected based on a lowest AIC value using bi-directional stepwise simplification (Table S1). In addition, paired t-tests were performed to further investigate the relationship between pre- and post-translocation measures of alpha diversity for each individually matched fish.

Microbiome structure (beta diversity), based on Bray-Curtis distance, was visualised using non-metric multidimensional scaling (NMDS) ordination, performed within mothur then plotted using ggplot2 (Wickham 2009). Multivariate statistical analysis of community separation (PERMANOVA) was performed using Adonis in the Vegan package (Oksanen et al. 2017). Initial (pre-translocation) differences in microbiome structure were investigated, including origin (wild/hatchery) and fish size (length) as fixed factors in the model. For the post-translocation sampling point, the effects of environment (hatchery/enriched/natural), origin, an environment:origin interaction and fish size on final microbiome structure were tested. The specific change in gut and skin community structure (Bray-Curtis distance) over time for individually-matched fish was also investigated using the model (ΔBC ∼ environment + origin + environment:origin + SGR). As before, the full models were reduced using stepwise simplification (Table S1).

Statistical analysis of OTU abundance was performed using DeSeq2 (Love et al. 2014), using rarefied data as recommended for microbiome libraries with large deviance in total library size between samples (Weiss et al. 2017). For the gut and skin separately, the effect of origin on initial (pre-translocation) OTU abundance was examined, while the effect of environment and origin on final (post-translocation) OTU abundance was identified using a multifactorial design including the main effects of environment, origin and their interaction. Within the DesSeq2 models, independent filtering of low coverage OTUS was applied, optimising power for identification of differentially abundant OTUs at a threshold of alpha=0.05. Default settings were applied for outlier detection and moderation of OTU level dispersion estimates. OTU abundance was considered significantly different at FDR <0.05.

## Results

### Specific growth rate and condition

There was a significant effect of both environment (hatchery, enriched, natural) and origin (hatchery or wild) on the specific growth rate for pre/post matched individual fish, and a significant interaction between factors (*Environment* F_2,42_ = 55.62, *P* < 0.001; *Origin* F_1,42_ = 53.18, *P* <0.001; *Environment:Origin* F_2,42_ = 5.02, *P* = 0.01; Figure S1). Specific growth rate was highest in the natural environment, and fish originating from the hatchery also showed a significantly higher growth rate than wild origin fish in both the hatchery and enriched groups. However, in the natural group there was no difference in growth rate between wild- and hatchery-origin fish. There was no significant effect of environment or origin on final condition index (*Environment* F_2,42_ = 2.41, *P* = 0.10; *Origin* F_1,42_ = 0.02, *P* = 0.88; *Environment:Origin* F_2,42_ = 2.10, *P* = 0.14).

### Microbial alpha diversity

Water microbial richness, but not diversity, was initially higher in the river compared to the hatchery (Chao1: t_2,3_ = 26.28, P=0.012; Shannon: t_1.03_ = 2.95, P=0.203), while at the second sampling point both water richness and diversity were significantly higher in the natural environment (leat) than in the hatchery and enriched tanks (Chao1: F_1.38_ = 11.41, P=0.025; Shannon: F_2,3_ = 18.67, P=0.020).

Gut microbial richness was initially significantly higher in wild than hatchery fish before the translocation, but there was no difference in gut Shannon diversity (Chao1: F_1,46_ = 62.85, P< 0.001; Shannon: F_1,46_ = 1.12, P = 0.295; Figure 2A). Overall, across all matched fish, gut microbial richness and diversity significantly declined during the course of the translocation experiment (Chao1: t_47_ = 5.40, P< 0.001; Shannon: t_47_ = 7.08, P<0.001; Figure 2C-H). In addition, the degree of individual-level change in gut Chao1 richness, but not Shannon diversity, was significantly affected by both environment and origin (Chao1: *Environment* F_2,44_ = 3.46, P=0.040; *Origin*-F_1,44_ = 41.90, P<0.001; Shannon: *Origin* F_1,46_ = 1.31, P = 0.259). Hatchery-origin fish fed an enriched diet showed a smaller reduction in gut microbial richness compared to those maintained in the same environment (hatchery to hatchery group), whilst those translocated to a natural environment actually experienced a small increase in gut richness. All wild-origin fish, which had higher initial gut richness, showed a greater decline in gut richness during the course of the translocation experiment compared to hatchery fish. Wild fish that experienced the largest change in environment and diet (wild to hatchery group) experienced a greater reduction in gut richness compared to those in the enriched and, especially, the natural environment groups. After the translocation, final gut Chao1 richness was strongly influenced by environment with no detectable effect of origin, size or the initial microbial richness for matched individuals (*Environment* F_2,44_ = 12.77, P< 0.001, *Length* F_1,44_ = 2.72, P = 0.10; Figure 2B). Fish maintained in natural conditions had higher gut richness than in both the hatchery and enriched groups, while an enriched diet also significantly increased gut Chao1 richness compared to that in the hatchery group. There was no significant effect of environment, origin or size on final gut Shannon diversity (*Environment* F_2,44_ = 1.77, P = 0.182, *Length* F_1,44_ = 3.18, P = 0.081).

**Figure 2.**
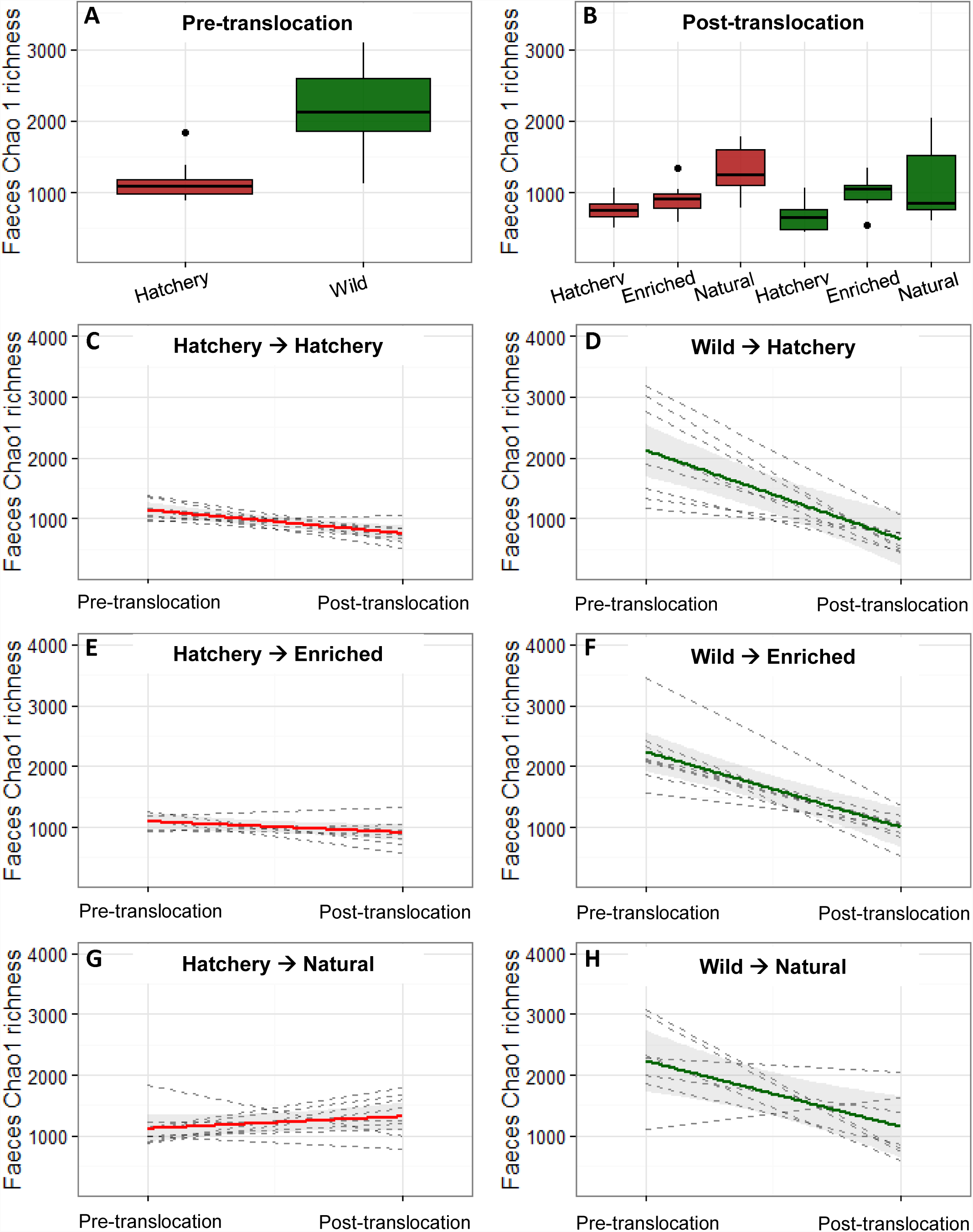
Faecal Chao1 richness in each group A) before (n=24) and B) after the experiment (n=8), red shading indicates hatchery origin and green shading indicates wild origin fish. C-H) Change in faecal Chao1 richness for all matched fish during the course of the experiment. Each dashed line represents an individual fish, with coloured lines displaying the average for each group and grey shading indicating 95% confidence intervals.

For the skin microbiome, there was no initial difference in richness or diversity between wild and hatchery fish (Chao1: *Origin* F_1,43_ = 3.64, P = 0.063, *Length* F_1,43_ = 0.19, P = 0.65, *Origin:Length* F_1,43_ = 2.72, P = 0.11; Shannon: *Origin* F_1,45_ = 0.54, P = 0.466). Across all fish, there was a small reduction in skin diversity but not richness over time (Chao1: t_46_ = −0.39, P = 0.70, Shannon: t_46_ = −2.26, P = 0.03), but environment had a significant effect on the degree of individual-level change in richness and diversity during the course of the translocation experiment (Chao1: *Environment* F_2,43_ = 4.38, P = 0.019; *Origin* F_1,43_ = 3.61, P=0.064; Shannon: *Environment* F_2,43_ = 7.64, P=0.001; *SGR* F_1,43_ = 3.28, P=0.077; Figure S2). Post translocation, there was also a significant effect of environment on final skin Chao1 richness and Shannon diversity (Chao1: *Environment* F_2,45_ = 4.55, P=0.016; Shannon: *Environment* F_2,43_ = 7.89, P = 0.001; *Pre-Shannon* F_1,43_ = 3.47, P = 0.07; Figure S2). In each case, fish from the natural environment had higher richness/diversity than in both the hatchery and enriched groups. There was no detectable effect of origin, size or the pre-translocation microbial richness/diversity for matched individuals on measures of final alpha diversity in any case.

### Microbial beta diversity

Before translocation, there was a significant difference in the structure of both the gut and the skin microbiome between wild and hatchery fish, and fish size also had a significant effect on beta diversity (Gut: *Origin* F_1,45_ = 3.35, P=0.016, *Length* F_1,45_ = 2.85, P=0.032; Skin: *Origin* F_1,44_ = 2.55, P=0.025, *Length* F_1,44_ = 2.15, P=0.050; Figure 3). Overall, during the course of the translocation experiment there was a large change in gut and skin microbiome structure across all fish, but environment and origin significantly affected the degree of structural change (Bray-Curtis distance for individually pre/post matched fish) that occurred for both the gut and skin microbiome (Gut: *Environment* F_2,42_ = 3.46, P=0.041; *Origin* F_1,42_ = 5.57, P=0.023; *Environment:Origin* F_2,42_ = 9.62, P<0.001; Skin: *Environment* F_2,43_ = 2.05, P=0.041, *Origin* F_1,43_ = 21.57, P<0.001; Figure S3). The smallest change in community structure occurred for hatchery fish maintained in the same conditions and fed the same diet, while the largest change was found in fish which experienced the greatest environmental and dietary change (i.e. wild-hatchery and hatchery-wild). After translocation, final skin and gut microbiome structure was strongly affected by environment as well as fish origin and fish size (Gut: *Environment* F_2,41_ = 6.41, P=0.001; *Origin* F_1,41_ = 2.07, P=0.029; *Length* F_1,41_ = 2.16, P=0.024; *Environment:Origin* F_2,41_ = 1.23, P=0.197; Skin: *Environment* F_2,40_ = 13.20, P=0.001; *Origin* F_1,40_ = 9.56, P=0.001; *Length* F_1,40_ = 0.49, P=0.774; *Environment:Origin* F_2,40_ = 0.97, P=0.427).

**Figure 3.**
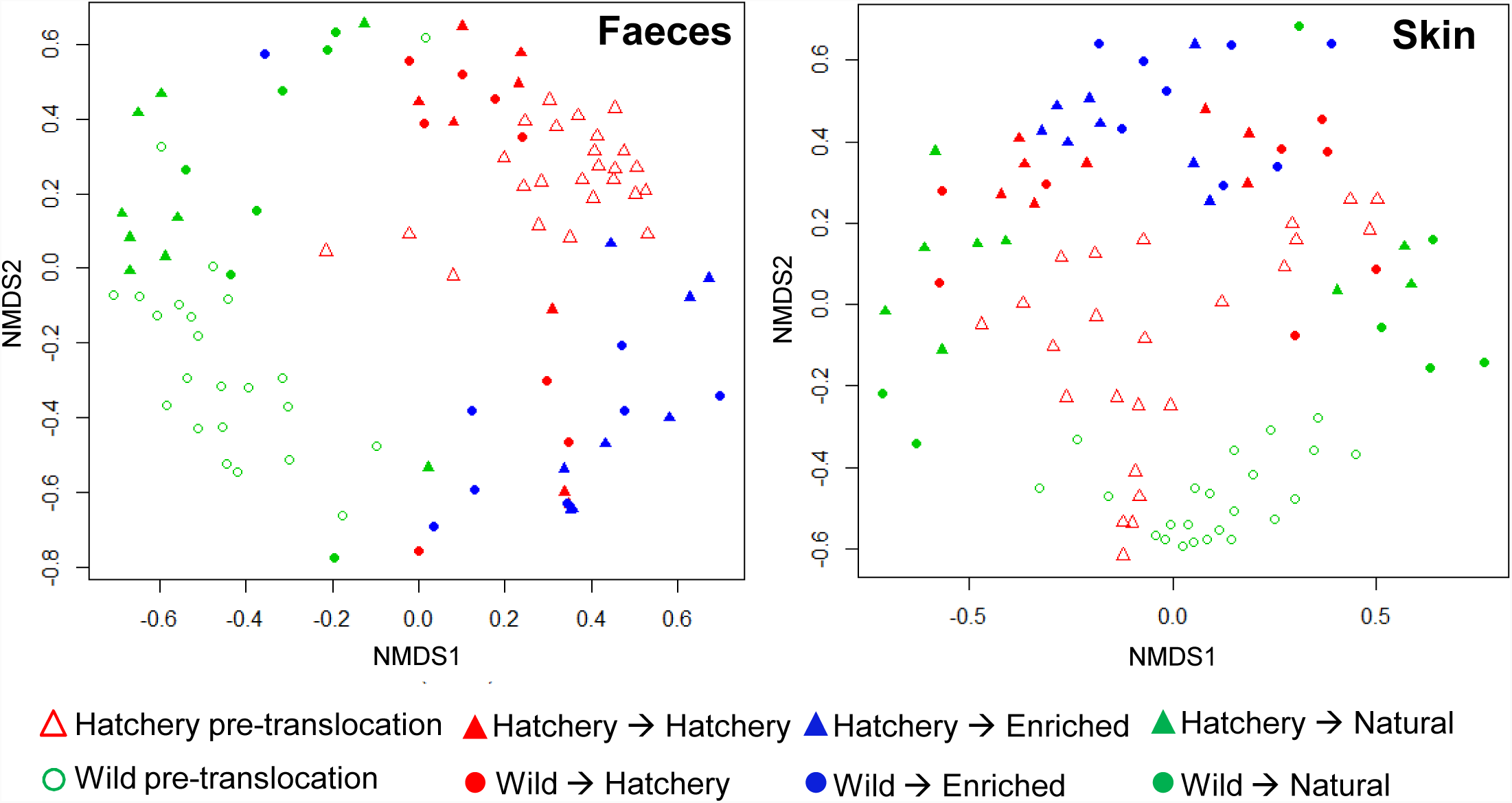
Non-metric multidimensional scaling (NMDS) ordination of microbial community structure based on Bray-Curtis distances, for all samples before and after the translocation experiment.

### OTU abundance

Before translocation, a total of 260 gut OTUs were significantly differentially abundant (FDR <0.05) between wild and hatchery fish (Table S2). The vast majority of them (236; 91%) were more abundant in wild fish but, notably, the OTUs which were more abundant in hatchery fish were predominantly Lactobacillales (14/24; 58%). Hatchery fish were dominated by several *Lactobacillus* sp., as well as an unclassified OTU from the order Bacillales. Wild fish were dominated by a similar unclassified OTU from the order Bacillales (93% sequence similarity to that in hatchery fish), as well as several OTUs from the family Enterobacteriaceae and from the Candidate division (CK.1C4.19) (Figure 4a, Figure S4a). In contrast, for the skin there were fewer initial differences in microbiome composition between wild and hatchery origin fish. Only 14 differentially abundant OTUs were identified, but these included an unclassified OTU from the family Rhodobiaceae which was amongst the most abundant OTUs in hatchery fish (Figure 4b, Figure S4b).

**Figure 4.**
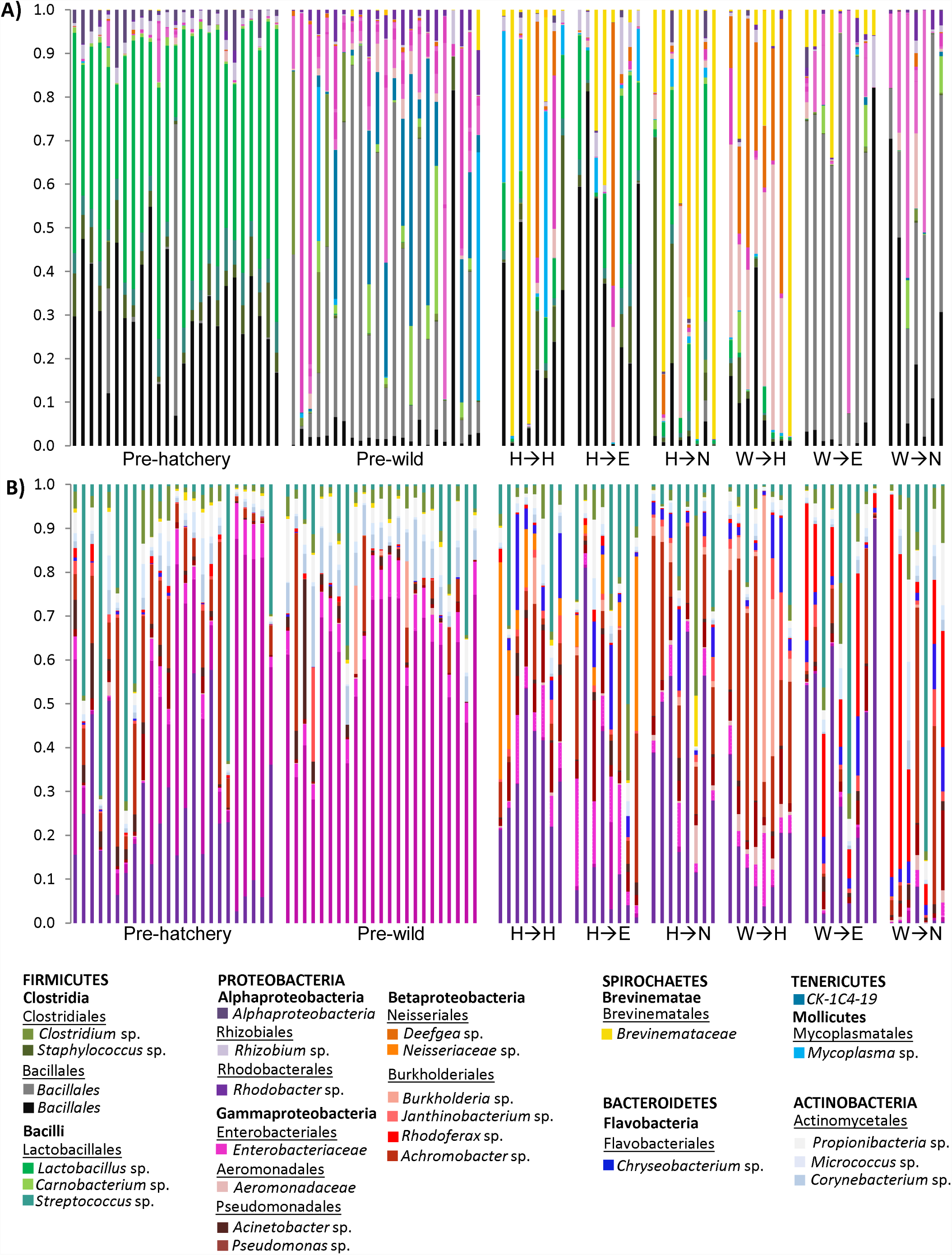
Relative abundance of the 25 most abundant OTUs in (A) faeces and (B) skin microbial communities before and after the translocation experiment. Each bar represents an individual fish. Origin; hatchery (H) and wild (W). Experimental environment; hatchery (H), enriched (E) and natural (N).

After translocation, in the gut a total of 88, 126 and 136 OTUs were differentially abundant between the hatchery-enriched, hatchery-natural and enriched-natural experimental groups, respectively (Table S2). In particular, the unclassified Bacillales OTU initially most abundant in wild fish was present at significantly higher levels in fish in the natural environment, while the phylogenetically similar unclassified Bacillales OTU initially dominant in hatchery fish was present at significantly higher levels in fish from both the hatchery and enriched environments. Additionally, fish from the hatchery environment harboured greater numbers of several *Lactobacillus* sp., *Mycoplasma* sp. and *Rhizobium* sp., while salmon fed the enriched diet displayed increased numbers of *Deefga* sp. and an OTU from the family Aeromonadaceae (Figure 4a, Figure S4a). In the natural environment group there were significantly lower levels of seven OTUs from the family Brevinemataceae and increased abundance of several OTUS from the family Enterobacteriaceae compared to fish from the hatchery and enriched groups. In addition to the effects of experimental environment, a total of 72 gut OTUs showed a significant effect of fish origin (Table S2). Thirty two OTUs from the family Brevinemataceae were all present at higher levels in hatchery-origin fish, together with some of the most dominant gut community members including *Mycoplasma* sp. and *Lactobacillus* sp., while OTUs more abundant in wild-origin fish included several OTUS from the family Enterobacteriaceae.

In the skin, a total of 14, 70 and 58 OTUs were differentially abundant between the hatchery-enriched, hatchery-natural and enriched-natural experimental groups, respectively (Figure 4b, Figure S4b, Table S3). Notably, the unclassified OTU from the family Rhodobiaceae, that was initially more abundant in hatchery fish before the translocation, was present at significantly higher levels in both the hatchery and enriched environment groups. This was also the only skin OTU significantly affected by fish origin, and was more abundant in hatchery-origin fish across all experimental environments.

### Community function

Before translocation, there were pronounced differences in predicted metagenomic function between wild and hatchery origin fish, in both the skin and the gut microbiome. In the gut there were 128 (39%) significantly differentially enriched KEGG pathways (FDR <0.05), with 188 (57%) in the skin. These included some of the most highly represented KEGG pathways such as ‘ABC Transporters’, ‘Purine metabolism’, ‘Two-component system’, and ‘Secretion System’.

After the translocation, in the gut 107 (32%) KEGG pathways were significantly differentially represented amongst experimental environments. Amongst the most represented KEGG pathways, these included ‘DNA repair and recombination proteins’, ‘Ribosome’, ‘Purine Metabolism’ and ‘Pyrimidine Metabolism’ (Figure S5a). In the skin microbiome, 103 (31%) KEGG pathways were differentially enriched amongst environmental treatments, including ‘Secretion system’ and ‘Replication, recombination and repair’ (Figure S5b). There was also a lasting effect of origin on seven (2.1%) gut KEGG pathways, and 57 (17 %) skin KEGG pathways (Figure S5).

## Discussion

Overall, we found evidence of extensive plasticity in the Atlantic salmon gut and skin microbiome associated with environmental change and development, but crucially we also identified some lasting effects of early life experience on microbial structure and function. Our results, demonstrating that both environmental plasticity and historical colonisation effects determine fish microbiome assemblage, are also likely to have important implications for host health and fitness.

### Change in the gut and skin microbiome over time

Using a powerful BACI approach, we identified a fundamental change in the gut microbiome over time. For each individually matched fish, initial gut alpha diversity had no detectable effect on final alpha diversity value after translocation and, overall, OTU richness and diversity both significantly declined over time. Gut community structure and function also clearly changed over time, including a general reduction in Actinobacteria OTUs and a notable increased abundance of OTUs from the family Brevinemataceae. These differences in diversity and structure are likely to reflect developmental changes in these juvenile salmon, as well as seasonal change across the six week experimental period. With maturation, the teleost gut microbiome tends to become less diverse and increasingly specialised and stable, reflecting a stronger influence of host-specific, microbial interactions and active dispersal (Burns et al. 2016; Stephens et al. 2016). For the skin, there were less pronounced changes in alpha diversity over time, but considerable changes in microbiome structure including a general reduction in Alphaproteobacteria, especially Enterobacteriaceae, and increased abundance of Betaproteobacteria, as well as Actinobacteria and Flavobacteria. These results suggest that temporal dynamics in the skin and gut microbiome differ. Little is known about developmental changes in the teleost skin microbiome, but seasonality is known to be an important determinant of fish and amphibian skin microbiota (Larsen et al. 2015; Longo et al. 2015).

### Microbiome plasticity in response to diet and environmental change

Before translocation, there was a clear distinction between the gut and skin microbiomes of hatchery- and wild-origin fish, consistent with our previous results showing differences in the diversity and structure of juvenile wild and hatchery-reared Atlantic salmon microbiomes (Uren Webster et al. 2018). Wild salmon had a richer and more diverse gut microbiome, with abundant Enterobacteriaceae compared to the dominance of Firmicutes (especially Lactobacillales) in hatchery fish, while the skin of wild and hatchery fish was dominated by different groups of Proteobacteria. However, following translocation, we found that microbiome structure, function and, especially, diversity was largely determined by environment, rather than fish origin and early-life experience. This suggests that, at a given time, the salmon microbiome is highly dependent on the current diet and environmental conditions that a fish is exposed to, and shows considerable plasticity following environmental change.

Salmon exposed to simulated wild conditions in the leat showed significantly higher microbial richness in the skin and, especially, the gut microbiome compared to that of fish in the hatchery environment. This is likely due to a higher degree of dietary diversity and variation compared to that of the artificial hatchery diet (Dehler et al. 2017; Orlov et al. 2006), with the higher growth rates observed under natural conditions also indicating that there was a seasonally rich and plentiful natural diet available to these fish. In addition, we found that microbial richness in the surrounding water was significantly higher in the leat than for the hatchery and enriched tanks, which is also likely to influence microbiome richness in both the gut and the skin (Boutin et al. 2013; Giatsis et al. 2015; Uren Webster et al. 2018). By enriching the standard hatchery diet with invertebrates, we were able to examine the effects of increased dietary diversity on the microbiome in isolation from other environmental variables. We found that gut and skin microbial richness increased with dietary enrichment, but to a much lesser extent than that observed in the natural environment which also encompassed wider environmental change. Furthermore, these pronounced effects of environment occurred regardless of fish origin, and despite the pronounced initial differences between wild and hatchery origin fish, demonstrating a capacity for extensive plasticity in microbial richness following environmental and dietary change.

Environment also had a marked effect on gut and skin microbiome structure, with a clear distinction between fish from all three experimental environments. This is consistent with a strong influence of diet (e.g. Gajardo et al. 2016a; Schmidt et al. 2016) as well as other environmental variables (e.g. Burns et al. 2016; Llewellyn et al. 2016; Smith et al. 2015) on gut microbiome structure in Atlantic salmon and other fish species. Evidence from fish in the enriched group shows that dietary change also specifically influences skin microbiome structure and function, although to a lesser extent than for the gut. Overall, our results highlight that there is considerable plasticity in the skin and gut microbiome structure in response to environmental change, regardless of fish origin and in spite of considerable initial differences between the wild and hatchery fish. Notably, fish subject to the largest change in diet and environmental conditions experienced the greatest change in microbiome structure, while hatchery fish maintained in hatchery conditions experienced the least.

Additionally, specific microbiome taxonomic composition was particularly distinct between fish in the natural experimental group, and those in the hatchery and enriched diet group, with a number of taxa that appeared to be distinctive of wild or hatchery conditions. In the gut we identified two highly abundant ‘wild-type’ and ‘hatchery-type’ OTUs within the order Bacillales (93% sequence similarity) that were consistently favoured by natural or hatchery conditions, before and after the translocation experiment. Multiple OTUs from the family Brevinemataceae became more abundant with time, but strikingly only in fish translocated to the hatchery and enriched environments. Brevinemataceae abundance is known to be variable in fish gut (Gajardo et al. 2016b; Pratte et al. 2018), and our results suggest that this taxa, besides being dependent on developmental stage, is strongly favoured by an artificial diet and/or other captive conditions and is signatory of hatchery environments. *Mycoplasma* sp. and several *Lactobacillus* sp. were also consistently enhanced in the hatchery-based environment, while several OTUs from family Enterobacteriaceae were much more abundant under natural conditions. This is consistent with what we found previously in wild and hatchery fish (Uren Webster et al. 2018), and highlights the potential for mutual exclusion between Enterobacteriaceae and *Lactobacillus* sp. similar to that in human gut (Sprockett et al. 2018).

Similarly, in the skin microbiome, we identified several OTUs that appeared to be specifically enhanced in captive conditions. These included an OTU from the family Rhodobiaceae, *Burkholderia* sp. and *Pseudomonas* sp., which were initially elevated in hatchery-origin fish and following translocation to the hatchery and enriched environments as well. However, compared to the gut, there appeared to be a less pronounced effect of experimental wild and hatchery conditions on the structural composition of skin microbial communities and fewer differentially abundant OTUs between groups. This could be because the skin microbiome is less strongly influenced by dietary change than the gut, and also likely reflects the fact that the hatchery/enriched groups were supplied with filtered river water with a similar microbial profile to that of the natural environment.

### Historical colonisation effects in the salmon microbiome

Fish origin had no discernable lasting effects on skin or gut microbial alpha diversity, which showed extensive plasticity and was apparently entirely determined by environment and developmental stage. However, we did identify persistent effects of fish origin on microbiome structure and function.

Hatchery conditions, in particular, left signatures in the gut and skin microbiome assembly regardless of environmental change, including the relative abundance of some of the most dominant community members, and in overall community structure. A number of OTUs were more abundant in hatchery-origin fish regardless of the environment they were translocated to. In the skin microbiome, a single very abundant OTU from the family Rhodobiaceae was initially far more prevalent, and remained more prevalent, in hatchery-origin fish translocated to each of the environmental groups. This OTU was also strongly enhanced by the hatchery and enriched conditions, but remained very rare in fish never exposed to hatchery conditions (wild to natural group). Several gut OTUs showed similar persistent patterns in hatchery fish, including some of the most dominant community members *Mycoplasma* sp. and *Lactobacillus* sp. However, the most striking effect of origin in the gut microbiome concerned the differential emergence of OTUs from the family Brevinemataceae. Although Brevinemataceae were rare before translocation, they became significantly more abundant in hatchery-origin fish across all experimental groups, even in fish translocated to natural conditions. This strongly suggests that early-life experience of captivity promotes Brevinemataceae emergence in the gut, regardless of later environmental conditions.

These effects of fish origin on community structure likely reflect differential microbial colonisation history. Priority effects describe how the order and timing of microbial arrival alters the availability of ecological niches for successive colonisers, through niche pre-emption or modification (Fukami 2015; Sprockett et al. 2018; Walter & Ley 2011). Fundamentally, within richer microbial communities, such as those found associated with the wild fish before translocation, higher competition is likely to reduce the chance of certain taxa becoming over-dominant (Fukami 2015; Hibbing et al. 2010). It could be that lasting historical signatures were more prevalent in hatchery-origin fish because reduced microbial competition allowed taxa favoured by hatchery conditions to flourish and establish dominance during early development. Large populations of established, dominant bacteria are likely to be able to outcompete later arrivals, and may also have an increased potential to adapt to environmental change, due to a higher probability of mutation and gene flow (Howe et al. 2015; Sprockett et al. 2018; Walter & Ley 2011). This may account for the continued presence of dominant OTUs, including *Mycoplasma* sp. and *Lactobacillus* sp., in hatchery-origin fish following translocation to the natural environment, but a lack of similar persistent, signatory OTUs in wild-origin fish. The preferential emergence of Brevinemataceae OTUs in hatchery-origin fish could be explained by niche modification, whereby previous colonisers in the hatchery gut provide favourable nutrients or conditions for Brevinemataceae growth, and/or conditions in the wild fish gut inhibit their successful colonisation (Fukami 2015).

In addition to priority effects due to microbial colonisation history, it is likely that host-specific differences between the wild and hatchery salmon populations may also contribute to the observed lasting signatures of origin on microbiome structure. These could include differences in the genetic background of the two fish populations. Host genotype is known to influence microbiome diversity and structure, although to a lesser extent than environmental factors, with the interaction between microbiota and the immune system likely to represent the predominant selective mechanism (Burns et al. 2017; Gensollen et al. 2016a; Stagaman et al. 2017; Uren Webster et al. 2018). Additionally, the two fish populations are likely to have experienced very different environmental conditions and pathogens in the wild and captivity, shaping the development of their adaptive immune system and, thus, the nature of selective processes influencing microbiota assembly (Foster et al. 2017; Gensollen et al. 2016a). Further research is required to establish the relative contributions of host genetic and environmentally-driven epigenetic influences on lasting signatures of origin in the fish microbiome, relative to historical microbial colonisation effects.

### Perspective

Overall, we show that the scope for environmental plasticity in the fish gut and skin microbiome is extensive, where degree of change in microbial community assembly closely mirrors the scale of environmental change. In addition, we also show, for the first time in fish, clear evidence that conditions experienced in early life can have a lasting influence on microbiome structure and function, likely primarily due to the influence of microbial colonisation history through priority effects. These results could have a range of important implications for evolutionary ecology, conservation and management. For example, extensive metagenomic plasticity could potentially contribute to host phenotypic plasticity, enhancing capacity to adapt to environmental challenges, such as climate change, emergent pathogens and pollution. Historical colonisation effects, potentially associated with an advantageous host phenotype in a given environment, could represent a mechanism contributing to local adaptation, or even phenotypic mismatch in hatchery-released fish used to supplement natural populations. In addition, our results demonstrate how common hatchery husbandry processes which alter microbial colonisation and succession, such as antimicrobial treatments, could have a lasting impact on the fish microbiome and, potentially, also on host health. However, they also highlight the possibility of conditioning the microbiome, for example through diet, to improve disease resistance in farmed fish or reduce phenotypic mismatch in hatchery released fish. Future research establishing causal links to host phenotype will be essential to establish the relative importance of microbiome plasticity and historical colonisation effects on host fitness.

## Acknowledgements

We are grateful to Paul Greest, Sophie Gott and Natural Resources Wales for providing us with access to the wild salmon, to John Taylor at the Cynrig fish culture unit for fish husbandry, to Matthew Hitchings for conducting the sequencing, and Chloe Robinson and Matteo Rolla for assistance with sampling. This work was funded by a BBSRC-NERC Aquaculture grant (BB/M026469/1) to CGL and the Welsh Government and Higher Education Funding Council for Wales (HEFCW) through the Sêr Cymru National Research Network for Low Carbon Energy and Environment (NRN-LCEE) to SC.

## Data Accessibility Statement

All Illumina sequence reads are will be available from the European Nucleotide Archive under the accession number PRJEB30953.

## Author Contributions

CGL, TUW, PG and SC designed the study; TUW, DRB and GC performed the experiment; TUW and DRB analysed the data; TUW, CGL and SC wrote the manuscript. All authors contributed to the final version of the manuscript.

## References

Alberdi, A., Aizpurua, O., Bohmann, K., Zepeda-Mendoza, M. L. & Gilbert, M. T. P. 2016 Do Vertebrate Gut Metagenomes Confer Rapid Ecological Adaptation? Trends in Ecology & Evolution 31, 689–699.

Bolger, A. M., Lohse, M. & Usadel, B. 2014 Trimmomatic: a flexible trimmer for Illumina sequence data. Bioinformatics 30, 2114–20.

Borre, Y. E., O’Keeffe, G. W., Clarke, G., Stanton, C., Dinan, T. G. & Cryan, J. F. 2014 Microbiota and neurodevelopmental windows: implications for brain disorders. Trends in Molecular Medicine 20, 509–518.

Boutin, S., Bernatchez, L., Audet, C. & Derôme, N. 2013 Network analysis highlights complex interactions between pathogen, host and commensal microbiota. 8, e84772.

Bray, J. R. & Curtis, J. T. 1957 An ordination of the upland forest communities of southern Wisconsin. Ecological monographs 27, 325–349.

Burns, A. R., Miller, E., Agarwal, M., Rolig, A. S., Milligan-Myhre, K., Seredick, S., Guillemin, K. & Bohannan, B. J. M. 2017 Interhost dispersal alters microbiome assembly and can overwhelm host innate immunity in an experimental zebrafish model. Proc Natl Acad Sci U S A 114, 11181–11186.

Burns, A. R., Stephens, W. Z., Stagaman, K., Wong, S., Rawls, J. F., Guillemin, K. & Bohannan, B. J. M. 2016 Contribution of neutral processes to the assembly of gut microbial communities in the zebrafish over host development. The ISME Journal 10, 655–664.

Caporaso, J. G., Kuczynski, J., Stombaugh, J., Bittinger, K., Bushman, F. D. & Costello, E. K. 2010 QIIME allows analysis of high-throughput community sequencing data. Nat Methods 7, 335–336.

Chen, L., Garmaeva, S., Zhernakova, A., Fu, J. & Wijmenga, C. 2018 A system biology perspective on environment–host–microbe interactions. Human Molecular Genetics 27, R187–R194.

Chevalier, C., Stojanovic, O., Colin, D. J., Suarez-Zamorano, N., Tarallo, V., Veyrat-Durebex, C., Rigo, D., Fabbiano, S., Stevanovic, A., Hagemann, S., Montet, X., Seimbille, Y., Zamboni, N., Hapfelmeier, S. & Trajkovski, M. 2015 Gut Microbiota Orchestrates Energy Homeostasis during Cold. Cell 163, 1360–74.

Davenport, E. R., Sanders, J. G., Song, S. J., Amato, K. R., Clark, A. G. & Knight, R. 2017 The human microbiome in evolution. BMC Biology 15, 127.

Dehler, C. E., Secombes, C. J. & Martin, S. A. 2017 Environmental and physiological factors shape the gut microbiota of Atlantic salmon parr (Salmo salar L.). Aquaculture 467, 149–157.

DeSantis, T. Z., Hugenholtz, P., Larsen, N., Rojas, M., Brodie, E. L., Keller, K., Huber, T., Dalevi, D., Hu, P. & Andersen, G. L. 2006 Greengenes, a chimera-checked 16S rRNA gene database and workbench compatible with ARB. Applied and environmental microbiology 72, 5069–5072.

Donnelly, W. & Whoriskey, F. 1993 Transplantation of Atlantic salmon (Salmo salar) and crypsis breakdown. Canadian Special Publication of Fisheries and Aquatic Sciences 118, 25–34.

Edgar, R. C., Haas, B. J., Clemente, J. C., Quince, C. & Knight, R. 2011 UCHIME improves sensitivity and speed of chimera detection. Bioinformatics 27, 2194–2200.

Foster, J. A., Rinaman, L. & Cryan, J. F. 2017 Stress & the gut-brain axis: Regulation by the microbiome. Neurobiology of Stress 19, 124–136.

Froese, R. 2006 Cube law, condition factor and weight–length relationships: history, meta-analysis and recommendations. Journal of Applied Ichthyology 22, 241–253.

Fukami, T. 2015 Historical contingency in community assembly: integrating niches, species pools, and priority effects. Annual Review of Ecology, Evolution, and Systematics 46, 1–23.

Gajardo, K., Jaramillo-Torres, A., Kortner, T. M., Merrifield, D. L., Tinsley, J., Bakke, A. M. & Krogdahl, Å. 2016a Alternative protein sources in the diet modulate microbiota and functionality in the distal intestine of Atlantic salmon (Salmo salar). Applied and environmental microbiology, AEM. 02615–16.

Gajardo, K., Rodiles, A., Kortner, T. M., Krogdahl, Å., Bakke, A. M., Merrifield, D. L. & Sørum, H. 2016b A high-resolution map of the gut microbiota in Atlantic salmon (Salmo salar): A basis for comparative gut microbial research. Scientific Reports 6, 30893.

Garcia de Leaniz, C., Fleming, I. A., Einum, S., Verspoor, E., Jordan, W. C., Consuegra, S., Aubin-Horth, N., Lajus, D., Letcher, B. H., Youngson, A. F., Webb, J. H., Vollestad, L. A., Villanueva, B., Ferguson, A. & Quinn, T. P. 2007 A critical review of adaptive genetic variation in Atlantic salmon: implications for conservation. Biol Rev Camb Philos Soc 82, 173–211.

Garcia de Leaniz, C., Fraser, N., Mikheev, V. & Huntingford, F. 1994 Individual recognition of juvenile salmonids using melanophore patterns. Journal of Fish Biology 45, 417–422.

Gensollen, T., Iyer, S. S., Kasper, D. L. & Blumberg, R. S. 2016a How colonization by microbiota in early life shapes the immune system. Science 352, 539–544.

Gensollen, T., Iyer, S. S., Kasper, D. L. & Blumberg, R. S. 2016b How colonization by microbiota in early life shapes the immune system. Science 352, 539–44.

Giatsis, C., Sipkema, D., Smidt, H., Heilig, H., Benvenuti, G., Verreth, J. & Verdegem, M. 2015 The impact of rearing environment on the development of gut microbiota in tilapia larvae. Scientific Reports 5, 18206.

Griffiths, A. M., Ellis, J. S., Clifton-Dey, D., Machado-Schiaffino, G., Bright, D., Garcia-Vazquez, E. & Stevens, J. R. 2011 Restoration versus recolonisation: The origin of Atlantic salmon (Salmo salar L.) currently in the River Thames. Biological Conservation 144, 2733–2738.

Hibbing, M. E., Fuqua, C., Parsek, M. R. & Peterson, S. B. 2010 Bacterial competition: surviving and thriving in the microbial jungle. Nature reviews. Microbiology 8, 15–25.

Hopkins, K. D. 1992 Reporting fish growth: a review of the basics. Journal of the world aquaculture society 23, 173–179.

Howe, A., Ringus, D. L., Williams, R. J., Choo, Z.-N., Greenwald, S. M., Owens, S. M., Coleman, M. L., Meyer, F. & Chang, E. B. 2015 Divergent responses of viral and bacterial communities in the gut microbiome to dietary disturbances in mice. The Isme Journal 10, 1217.

Korpela, K., Costea, P., Coelho, L. P., Kandels-Lewis, S., Willemsen, G., Boomsma, D. I., Segata, N. & Bork, P. 2018 Selective maternal seeding and environment shape the human gut microbiome. Genome research 28, 561–568.

Koskella, B., Hall, L. J. & Metcalf, C. J. E. 2017 The microbiome beyond the horizon of ecological and evolutionary theory. Nature ecology & evolution 1, 1606–1016.

Kozich, J. J., Westcott, S. L., Baxter, N. T., Highlander, S. K. & Schloss, P. D. 2013 Development of a dual-index sequencing strategy and curation pipeline for analyzing amplicon sequence data on the MiSeq Illumina sequencing platform. Appl Environ Microbiol 79, 5112–20.

Langille, M. G., Zaneveld, J., Caporaso, J. G., McDonald, D., Knights, D., Reyes, J. A., Clemente, J. C., Burkepile, D. E., Thurber, R. L. V. & Knight, R. 2013 Predictive functional profiling of microbial communities using 16S rRNA marker gene sequences. Nature biotechnology 31, 814.

Larsen, A. M., Bullard, S. A., Womble, M. & Arias, C. R. 2015 Community structure of skin microbiome of gulf killifish, Fundulus grandis, is driven by seasonality and not exposure to oiled sediments in a Louisiana salt marsh. Microbial ecology 70, 534–544.

Llewellyn, M. S., McGinnity, P., Dionne, M., Letourneau, J., Thonier, F., Carvalho, G. R., Creer, S. & Derome, N. 2016 The biogeography of the atlantic salmon (Salmo salar) gut microbiome. Isme J 10, 1280–4.

Lokesh, J. & Kiron, V. 2016 Transition from freshwater to seawater reshapes the skin-associated microbiota of Atlantic salmon. Scientific Reports 6, 19707.

Longo, A. V., Savage, A. E., Hewson, I. & Zamudio, K. R. 2015 Seasonal and ontogenetic variation of skin microbial communities and relationships to natural disease dynamics in declining amphibians. Royal Society open science 2, 140377.

Love, M. I., Huber, W. & Anders, S. 2014 Moderated estimation of fold change and dispersion for RNA-seq data with DESeq2. Genome Biology 15, 550–555.

Macke, E., Callens, M., De Meester, L. & Decaestecker, E. 2017 Host-genotype dependent gut microbiota drives zooplankton tolerance to toxic cyanobacteria. Nature Communications 8, 1608.

McDonald, D., Clemente, J. C., Kuczynski, J., Rideout, J. R., Stombaugh, J., Wendel, D., Wilke, A., Huse, S., Hufnagle, J., Meyer, F., Knight, R. & Caporaso, J. G. 2012 The Biological Observation Matrix (BIOM) format or: how I learned to stop worrying and love the ome-ome. GigaScience 1, 7.

Oksanen, J., Blanchet, F. G., Friendly, M., Kindt, R., Legendre, P., McGlinn, D., Minchin, P. R., O’Hara, R. B., Simpson, G. L., Solymos, P., Stevens, M. H. H., Szoecs, E. & Wagner, H. 2017 vegan: Community Ecology Package: https://CRAN.R-project.org/package=vegan.

Orlov, A. V., Gerasimov, Y. V. & Lapshin, O. M. 2006 The feeding behaviour of cultured and wild Atlantic salmon, Salmo salar L., in the Louvenga River, Kola Peninsula, Russia. ICES Journal of Marine Science: Journal du Conseil 63, 1297–1303.

Parks, D. H. & Beiko, R. G. 2010 Identifying biologically relevant differences between metagenomic communities. Bioinformatics 26, 715–21.

Pratte, Z. A., Besson, M., Hollman, R. D. & Stewart, F. J. 2018 The gills of reef fish support a distinct microbiome influenced by host-specific factors. Applied and environmental microbiology, AEM. 00063–18.

Quast, C., Pruesse, E., Yilmaz, P., Gerken, J., Schweer, T., Yarza, P., Peplies, J. & Glöckner, F. O. 2013 The SILVA ribosomal RNA gene database project: improved data processing and web-based tools. Nucleic Acids Research 41, D590–D596.

R_Core_Team. 2014 R: A language and environment for statistical computing. Vienna, Austria.: R Foundation for Statistical Computing.

Roberts, L. J., Taylor, J., Gough, P. J., Forman, D. W. & Garcia de Leaniz, C. 2014 Silver spoons in the rough: can environmental enrichment improve survival of hatchery Atlantic salmon Salmo salar in the wild? Journal of Fish Biology 85, 1972–1991.

Schmidt, V., Amaral-Zettler, L., Davidson, J., Summerfelt, S. & Good, C. 2016 The influence of fishmeal-free diets on microbial communities in Atlantic salmon Salmo salar recirculation aquaculture systems. Applied and environmental microbiology, AEM. 00902–16.

Smith, C. C., Snowberg, L. K., Caporaso, J. G., Knight, R. & Bolnick, D. I. 2015 Dietary input of microbes and host genetic variation shape among-population differences in stickleback gut microbiota. ISME J 9, 2515–2526.

Sprockett, D., Fukami, T. & Relman, D. A. 2018 Role of priority effects in the early-life assembly of the gut microbiota. Nature Reviews Gastroenterology & Hepatology 15, 197.

Stagaman, K., Burns, A. R., Guillemin, K. & Bohannan, B. J. 2017 The role of adaptive immunity as an ecological filter on the gut microbiota in zebrafish. Isme j 11, 1630–1639.

Stephens, W. Z., Burns, A. R., Stagaman, K., Wong, S., Rawls, J. F., Guillemin, K. & Bohannan, B. J. 2016 The composition of the zebrafish intestinal microbial community varies across development. Isme j 10, 644–54.

Suzuki, T. A. 2017 Links between Natural Variation in the Microbiome and Host Fitness in Wild Mammals. Integrative and comparative biology 57, 756–769.

Uren Webster, T. M., Consuegra, S., Hitchings, M. & Garcia de Leaniz, C. 2018 Inter-population variation in the Atlantic salmon microbiome reflects environmental and genetic diversity. Appl Environ Microbiol 84, e0061–18.

Vellend, M. 2016 The theory of ecological communities (MPB-57): Princeton University Press.

Walter, J. & Ley, R. 2011 The human gut microbiome: ecology and recent evolutionary changes. Annual review of microbiology 65, 411–429.

Weiss, S., Xu, Z. Z., Peddada, S., Amir, A., Bittinger, K., Gonzalez, A., Lozupone, C., Zaneveld, J. R., Vázquez-Baeza, Y., Birmingham, A., Hyde, E. R. & Knight, R. 2017 Normalization and microbial differential abundance strategies depend upon data characteristics. Microbiome 5, 27.

Wickham, H. 2009 ggplot2: Elegant Graphics for Data Analysis: Springer-Verlag New York.

Yassour, M., Vatanen, T., Siljander, H., Hamalainen, A. M., Harkonen, T., Ryhanen, S. J., Franzosa, E. A., Vlamakis, H., Huttenhower, C., Gevers, D., Lander, E. S., Knip, M. & Xavier, R. J. 2016 Natural history of the infant gut microbiome and impact of antibiotic treatment on bacterial strain diversity and stability. Sci Transl Med 8, 343ra81.

Zarkasi, K. Z., Abell, G. C., Taylor, R. S., Neuman, C., Hatje, E., Tamplin, M. L., Katouli, M. & Bowman, J. P. 2014 Pyrosequencing-based characterization of gastrointestinal bacteria of Atlantic salmon (Salmo salar L.) within a commercial mariculture system. J Appl Microbiol 117, 18–27.

