## Supporting Information for "Environmental plasticity and colonisation history in the Atlantic salmon microbiome: a translocation experiment"

**This supporting information contains:**

**P2-3: Table S1.** Selection of linear models based on AIC values.

**P4-9: Table S2.** Differentially abundant gut OTUs.

**P10-12: Table S3.** Differentially abundant skin OTUs.

**P13: Figure S1.** Specific growth rates for individual fish during the experiment.

**P14: Figure S2.** The effect of treatment and origin on skin Chao1 richness.

**P15: Figure S3.** The effect of treatment and origin on the change in microbiome structure.

**P16: Figure S4.** Heatmap illustrating the effect of treatment and origin on OTU abundance.

**P17: Figure S5.** Functional enrichment analysis.

**Table S1. Model selection table. Selection of linear models, based on AIC (Akaike Information Criterion) values, using the *Step* function. Highlighted models indicates selected optimal model.**

| | Model | AIC | ΔAIC |
|---|---|---|---|
| | SGR ~ environment + origin + environment:origin | -208.50 | 0.00 |
| | k ~ environment + origin + environment:origin | -216.55 | 0.00 |
| | k ~ environment + origin | -215.98 | 0.57 |
| Pre-translocation alpha diversity | F_chao_pre ~ origin | 593.18 | 0.00 |
| | F_chao_pre ~ origin + pre.length | 595.09 | -1.91 |
| | F_chao_pre ~origin + pre.length + origin:pre.length | 596.87 | -3.69 |
| | F_shannon_pre ~ origin | -37.72 | 0.00 |
| | F_shannon_pre ~ pre.length | -36.85 | -0.87 |
| | F_shannon_pre ~ origin + pre.length | -35.72 | -2.00 |
| | F_shannon_pre ~ origin + pre.length + origin:pre.length | -34.13 | -3.59 |
| | S_chao_pre ~ origin + pre.length + origin:pre.length | 489.44 | 0.00 |
| | S_shannon_pre ~ origin | -70.94 | 0.00 |
| | S_shannon_pre ~ pre.length | -70.82 | -0.12 |
| | S_shannon_pre ~ origin + pre.length | -70.23 | -0.71 |
| | S_shannon_pre ~ origin + pre.length + origin:pre.length | -69.72 | -1.22 |
| Post-translocation alpha diversity | F_chao_post ~ environment + post_length | 550.56 | 0.00 |
| | F_chao_post ~ environment + F_chao_pre + post_length | 550.83 | -0.27 |
| | F_chao_post ~ environment | 551.44 | -0.88 |
| | F_chao_post ~ environment + origin + post_length | 551.96 | -1.40 |
| | F_chao_post ~ environment + F_chao_pre | 552.09 | -1.53 |
| | F_chao_post ~ environment + origin + F_chao_pre + post_length | 552.72 | -2.16 |
| | F_chao_post ~ environment + origin + F_chao_pre | 553.96 | -3.40 |
| | F_shannon_post ~ environment + post_length | -5.38 | 0.00 |
| | F_shannon_post ~ post.length | -4.56 | -0.83 |
| | F_shannon_post ~ environment + F_shannon_pre + post_length | -3.96 | -1.42 |
| | F_shannon_post ~ environment + origin + post_length | -3.73 | -1.65 |
| | F_shannon_post ~ environment | -3.21 | -2.17 |
| | F_shannon_post ~ environment + origin + F_shannon_pre + post_length | -2.20 | -3.18 |
| | F_shannon_post ~ environment + F_shannon_pre | -2.12 | -3.26 |
| | F_shannon_post ~ origin + F_shannon_pre + post_length | -1.40 | -3.98 |
| | S_chao_post ~ environment | 494.31 | 0.00 |
| | S_chao_post ~ environment + origin | 495.25 | -0.94 |
| | S_chao_post ~ environment + S_chao_pre | 495.26 | -0.95 |
| | S_chao_post ~ environment + origin + S_chao_pre | 496.65 | -2.34 |
| | S_chao_post ~ environment + S_chao_pre + post_length | 496.68 | -2.37 |
| | S_chao_post ~ environment + origin + post_length | 496.96 | -2.65 |
| | S_chao_post ~ environment + origin + S_chao_pre + post_length | 498.07 | -3.76 |
| | S_chao_post ~ origin + S_chao_pre + post_length | 498.24 | -3.93 |
| | S_shannon_post ~ environment + S_shannon_pre | -64.63 | 0.00 |
| | S_shannon_post ~ environment + origin + S_shannon_pre | -64.13 | -0.49 |
| | S_shannon_post ~ environment + S_shannon_pre + post_length | -63.07 | -1.55 |
| | S_shannon_post ~ environment | -62.98 | -1.65 |
| | S_shannon_post ~ environment + origin | -62.96 | -1.66 |
| | S_shannon_post ~ environment + S_shannon_pre + post_length | -62.53 | -2.10 |
| | S_shannon_post ~ environment + origin + post_length | -62.11 | -2.52 |
| | S_shannon_post ~ environment + origin + environment:origin + S_shannon_pre | -61.54 | -3.09 |
| Change in α diversity | DF_Chao ~ environment + origin | 619.23 | 0.00 |
| | DF_Chao ~ origin | 620.57 | -1.34 |
| | DF_Chao ~ origin + SGR | 620.79 | -1.56 |
| | DF_Chao ~ environment + origin + SGR | 621.15 | -1.92 |
| | DF_Chao ~ environment + origin + environment:origin | 622.87 | -3.64 |
| | DF_Shannon ~ environment + origin | 16.56 | 0.00 |
| | DF_Shannon ~ origin + SGR | 18.56 | -2.00 |
| | DF_Shannon ~ environment + SGR | 20.39 | -3.82 |

| | | | |
|---|---|---|---|
| | DF_Shannon ~ origin | 20.42 | -3.85 |
| | DS_Chao ~ environment + SGR | 530.50 | 0.00 |
| | DS_Chao ~ environment + origin + SGR | 532.34 | -1.84 |
| | DS_Chao ~ environment + origin | 534.18 | -3.68 |
| | DS_Shannon ~ environment + SGR | -25.40 | 0.00 |
| | DS_Shannon ~ environment + origin | -24.03 | -1.37 |
| | DS_Shannon ~ environment | -23.95 | -1.45 |
| | DS_Shannon ~ environment + origin + SGR | -23.43 | -1.96 |
| | DS_Shannon ~ environment + origin + environment:origin | -23.13 | -2.27 |
| | DS_Shannon ~ environment + origin + environment:origin + SGR | -21.40 | -4.00 |
| **Change in β diversity** | F_BC_distance ~ environment + origin + environment:origin | -213.21 | 0.00 |
| | F_BC_distance ~ environment + origin + environment:origin + SGR | -211.70 | -1.51 |
| | S_BC_distance ~ environment + origin | -214.28 | 0.00 |
| | S_BC_distance ~ origin | -213.44 | -0.84 |
| | S_BC_distance ~ origin + SGR | -213.27 | -1.01 |
| | S_BC_distance ~ environment + origin + SGR | -212.34 | -1.94 |
| | S_BC_distance ~ environment + origin + environment:origin | -211.26 | -3.02 |

**Table S2. Differentially abundant gut OTUs.**

| OTU | Annotation | H:N Log2 Fold Change | H:W FDR | H:E Log2 Fold Change | H:E FDR | E:N Log2 Fold Change | E:W FDR | H:W origin Log2 Fold Change | Origin FDR |
|---|---|---|---|---|---|---|---|---|---|
| Otu00001 | Bacteria_unclassified | 0.00 | 1.000 | 0.00 | 1.000 | 0.00 | 1.000 | 0.00 | 1.000 |
| Otu00002 | Alphaproteobacteria_unclassified | -1.48 | 0.451 | -3.21 | 0.100 | 1.73 | 0.321 | 3.49 | 0.034 |
| Otu00003 | Bacteria_unclassified | -11.64 | 2.47E-27 | -2.51 | 0.070 | -9.13 | 4.73E-16 | 5.03 | 9.75E-07 |
| Otu00004 | Enterobacteriaceae_unclassified | -9.75 | 1.62E-13 | -1.08 | 0.478 | -8.67 | 2.56E-09 | -2.93 | 0.039 |
| Otu00006 | Brevinemataceae_unclassified | 6.46 | 1.85E-04 | -2.97 | 0.124 | 9.42 | 1.32E-08 | 7.39 | 8.73E-07 |
| Otu00007 | Lactobacillus | 9.47 | 7.91E-18 | -2.16 | 0.028 | 11.63 | 2.09E-27 | 1.96 | 0.053 |
| Otu00008 | Achromobacter | 2.36 | 0.256 | -5.21 | 0.005 | 7.57 | 6.37E-05 | 2.76 | 0.128 |
| Otu00010 | Aeromonadaceae_unclassified | 1.76 | 0.263 | -3.53 | 0.027 | 5.28 | 3.13E-04 | -2.47 | 0.071 |
| Otu00011 | Propionibacterium | -0.25 | 0.787 | -2.57 | 0.002 | 2.31 | 0.005 | 2.24 | 0.003 |
| Otu00012 | Mycoplasma | 7.55 | 5.53E-07 | 1.10 | 0.448 | 6.45 | 2.33E-05 | 4.84 | 4.96E-04 |
| Otu00013 | Staphylococcus | 6.05 | 3.24E-07 | -3.82 | 6.10E-04 | 9.86 | 3.16E-18 | 3.86 | 3.41E-04 |
| Otu00014 | Staphylococcus | 0.08 | 0.941 | -3.24 | 3.08E-04 | 3.33 | 1.54E-04 | 2.07 | 0.015 |
| Otu00015 | Enterobacteriaceae_unclassified | 4.30 | 9.86E-04 | 1.01 | 0.448 | 3.29 | 0.016 | -0.78 | 0.534 |
| Otu00016 | Ruminococcaceae_unclassified | -9.76 | 2.79E-07 | -3.78 | 0.096 | -5.98 | 0.003 | 3.91 | 0.036 |
| Otu00017 | CK | -8.32 | 1.16E-05 | -2.57 | 0.291 | -5.75 | 0.005 | 4.76 | 0.015 |
| Otu00018 | Rhodoferax | -4.95 | 5.30E-04 | -2.91 | 0.075 | -2.03 | 0.168 | 2.05 | 0.140 |
| Otu00020 | Corynebacterium | -0.34 | 0.847 | -3.56 | 0.021 | 3.22 | 0.040 | 3.91 | 0.005 |
| Otu00021 | Clostridium | -7.26 | 0.001 | -5.19 | 0.043 | -2.07 | 0.342 | 5.66 | 0.010 |
| Otu00022 | Streptococcus | 4.54 | 6.69E-06 | -2.26 | 0.026 | 6.80 | 2.11E-12 | 2.86 | 0.002 |
| Otu00023 | Chryseobacterium | 0.19 | 0.899 | -1.66 | 0.267 | 1.85 | 0.172 | 2.95 | 0.017 |
| Otu00024 | Pseudomonas | -1.40 | 0.354 | -4.95 | 3.08E-04 | 3.55 | 0.012 | -0.46 | 0.741 |
| Otu00025 | Acinetobacter | -0.22 | 0.874 | -4.51 | 2.07E-04 | 4.29 | 3.26E-04 | 1.89 | 0.095 |
| Otu00026 | Deefgea | 7.43 | 3.38E-06 | -0.33 | 0.830 | 7.76 | 1.06E-06 | -1.96 | 0.235 |
| Otu00027 | Micrococcus | 0.62 | 0.613 | -2.70 | 0.023 | 3.32 | 0.004 | 1.76 | 0.095 |
| Otu00028 | Enhydrobacter | 3.50 | 4.91E-04 | -2.53 | 0.008 | 6.03 | 2.12E-10 | 2.83 | 0.002 |
| Otu00033 | Lactobacillus | 6.48 | 3.24E-07 | -2.13 | 0.052 | 8.61 | 2.35E-12 | 3.49 | 0.002 |
| Otu00034 | Deefgea | -1.80 | 0.303 | -6.91 | 1.91E-05 | 5.11 | 0.002 | -1.63 | 0.326 |
| Otu00035 | Carnobacterium | -7.18 | 4.52E-10 | -5.47 | 9.30E-06 | -1.71 | 0.163 | 0.50 | 0.679 |
| Otu00036 | Rhodococcus | -0.20 | 0.874 | -2.25 | 0.068 | 2.05 | 0.085 | 0.91 | 0.413 |
| Otu00038 | Brevinemataceae_unclassified | 6.21 | 0.003 | -3.04 | 0.175 | 9.26 | 4.03E-06 | 7.35 | 5.52E-05 |
| Otu00039 | Rhizobium | 0.24 | 0.790 | 0.20 | 0.789 | 0.04 | 0.953 | 0.69 | 0.391 |
| Otu00040 | Janthinobacterium | -1.52 | 0.237 | -3.75 | 0.003 | 2.23 | 0.079 | 2.13 | 0.058 |
| Otu00042 | Dermacoccus | 5.10 | 4.96E-05 | -2.27 | 0.034 | 7.38 | 1.21E-09 | 3.65 | 0.001 |
| Otu00044 | Sediminibacterium | -0.59 | 0.569 | -2.07 | 0.042 | 1.48 | 0.130 | 1.58 | 0.070 |
| Otu00045 | Rhodobacter | -9.05 | 3.88E-14 | -6.23 | 2.03E-06 | -2.82 | 0.031 | -0.06 | 0.965 |

| | | | | | | | | | |
|---|---|---|---|---|---|---|---|---|---|
| Otu00046 | Enterobacter | 2.46 | 0.096 | -1.36 | 0.356 | 3.82 | 0.008 | 0.93 | 0.485 |
| Otu00047 | Delftia | -1.01 | 0.544 | -2.50 | 0.126 | 1.50 | 0.312 | 2.66 | 0.052 |
| Otu00048 | Pseudomonas | -0.20 | 0.874 | -4.77 | 4.38E-06 | 4.58 | 9.24E-06 | 1.01 | 0.364 |
| Otu00049 | Rickettsiella | -9.59 | 4.15E-08 | -14.12 | 7.58E-17 | 4.52 | 5.80E-04 | 0.09 | 0.961 |
| Otu00051 | Proteobacteria_unclassified | -2.29 | 0.420 | -6.26 | 0.020 | 3.98 | 0.127 | 2.19 | 0.396 |
| Otu00052 | Sphingomonas | -1.56 | 0.196 | -2.33 | 0.062 | 0.76 | 0.487 | 2.04 | 0.053 |
| Otu00053 | Rhodococcus | 0.66 | 0.562 | -4.22 | 1.04E-05 | 4.89 | 5.01E-07 | 2.78 | 0.003 |
| Otu00054 | Roseomonas | -9.77 | 2.89E-12 | -0.79 | 0.623 | -8.98 | 8.95E-09 | 0.86 | 0.589 |
| Otu00056 | Comamonadaceae_unclassified | -2.58 | 0.049 | -1.90 | 0.191 | -0.68 | 0.573 | 1.63 | 0.196 |
| Otu00059 | Pseudomonas | -1.01 | 0.396 | -5.41 | 9.38E-08 | 4.41 | 1.60E-05 | 1.21 | 0.255 |
| Otu00060 | Variovorax | -1.44 | 0.147 | -3.39 | 3.06E-04 | 1.95 | 0.045 | 1.96 | 0.024 |
| Otu00061 | Betaproteobacteria_unclassified | -3.01 | 0.201 | -11.68 | 3.49E-08 | 8.67 | 2.55E-05 | 0.29 | 0.889 |
| Otu00065 | Bradyrhizobium | -1.61 | 0.083 | -2.15 | 0.025 | 0.55 | 0.522 | 1.09 | 0.219 |
| Otu00066 | Enterobacteriaceae_unclassified | -7.29 | 3.79E-05 | -7.07 | 1.56E-04 | -0.23 | 0.895 | 2.22 | 0.219 |
| Otu00067 | Lactobacillus | 7.87 | 4.17E-09 | -2.13 | 0.033 | 10.00 | 1.40E-14 | 2.70 | 0.019 |
| Otu00068 | Rhodobacter | -6.24 | 1.45E-05 | -6.84 | 7.10E-06 | 0.59 | 0.674 | -0.06 | 0.969 |
| Otu00071 | Weissella | 8.27 | 4.17E-09 | -2.31 | 0.037 | 10.58 | 1.03E-14 | 2.64 | 0.030 |
| Otu00072 | Leuconostoc | 8.35 | 1.70E-10 | -2.42 | 0.010 | 10.77 | 2.99E-17 | 2.52 | 0.026 |
| Otu00078 | Pseudomonas | -3.08 | 0.037 | -7.58 | 3.49E-08 | 4.50 | 0.001 | 3.04 | 0.019 |
| Otu00079 | Sphingobacterium | 0.59 | 0.772 | -2.69 | 0.145 | 3.28 | 0.078 | 3.90 | 0.016 |
| Otu00082 | Flavobacterium | -2.18 | 0.173 | -2.19 | 0.198 | 0.01 | 0.995 | 2.51 | 0.071 |
| Otu00083 | Aeromonadaceae_unclassified | 0.69 | 0.784 | -12.81 | 3.53E-13 | 13.50 | 2.11E-12 | 1.25 | 0.515 |
| Otu00084 | Bacteria_unclassified | -14.80 | 7.18E-17 | -2.08 | 0.399 | -12.72 | 1.24E-11 | 2.87 | 0.143 |
| Otu00085 | Limnohabitans | -9.00 | 3.35E-04 | -3.31 | 0.295 | -5.69 | 0.032 | 3.34 | 0.196 |
| Otu00087 | Vagococcus | 5.56 | 1.42E-04 | -2.48 | 0.052 | 8.04 | 9.73E-09 | 1.25 | 0.382 |
| Otu00088 | Paracoccus | 1.18 | 0.454 | -5.13 | 1.47E-04 | 6.31 | 3.04E-06 | 0.97 | 0.468 |
| Otu00090 | Lactobacillus | 0.59 | 0.689 | -3.60 | 0.003 | 4.19 | 0.001 | 0.97 | 0.436 |
| Otu00091 | Bacteria_unclassified | -12.90 | 1.07E-16 | -4.04 | 0.052 | -8.86 | 4.55E-09 | 4.15 | 0.012 |
| Otu00093 | Bosea | 0.12 | 0.906 | -0.16 | 0.851 | 0.28 | 0.750 | 0.50 | 0.570 |
| Otu00095 | Actinomycetales_unclassified | -4.40 | 0.009 | -4.70 | 0.007 | 0.30 | 0.849 | 0.78 | 0.628 |
| Otu00101 | Actinomycetales_unclassified | -6.77 | 1.03E-04 | -9.55 | 3.49E-08 | 2.78 | 0.087 | 3.52 | 0.026 |
| Otu00102 | Agrobacterium | -1.70 | 0.366 | -4.28 | 0.020 | 2.58 | 0.144 | 0.27 | 0.871 |
| Otu00103 | Clostridiales_unclassified | -9.34 | 9.39E-08 | -5.56 | 0.004 | -3.78 | 0.043 | 0.20 | 0.909 |
| Otu00105 | Methylobacterium | 0.76 | 0.601 | -1.36 | 0.351 | 2.12 | 0.124 | 2.97 | 0.017 |
| Otu00108 | Lactobacillus | 7.21 | 1.91E-07 | -3.08 | 0.002 | 10.30 | 1.03E-14 | 2.36 | 0.047 |
| Otu00109 | Flavobacterium | -0.53 | 0.777 | -3.86 | 0.024 | 3.33 | 0.046 | 0.87 | 0.577 |
| Otu00117 | Lactobacillus | 4.97 | 2.62E-04 | -2.51 | 0.037 | 7.48 | 9.32E-09 | 4.28 | 2.85E-04 |
| Otu00118 | Oxalobacteraceae_unclassified | -7.08 | 4.58E-06 | -6.32 | 1.35E-04 | -0.76 | 0.602 | 0.79 | 0.610 |
| Otu00122 | Microbacteriaceae_unclassified | -3.20 | 0.002 | -3.57 | 7.29E-04 | 0.38 | 0.701 | 0.55 | 0.582 |

| | | | | | | | | | |
|---|---|---|---|---|---|---|---|---|---|
| Otu00125 | Bacteria_unclassified | -0.96 | 0.066 | -0.44 | 0.426 | -0.53 | 0.326 | 0.01 | 0.979 |
| Otu00126 | Clostridium | -3.06 | 0.451 | -11.02 | 0.003 | 7.96 | 0.028 | -2.52 | 0.468 |
| Otu00127 | Caulobacteraceae_unclassified | -1.49 | 0.391 | -3.83 | 0.018 | 2.34 | 0.145 | 2.95 | 0.041 |
| Otu00129 | Flectobacillus | -6.64 | 7.03E-04 | -6.59 | 0.001 | -0.05 | 0.977 | 1.09 | 0.565 |
| Otu00131 | Novosphingobium | -5.14 | 0.001 | -2.68 | 0.169 | -2.46 | 0.154 | 3.97 | 0.013 |
| Otu00134 | Massilia | -1.50 | 0.508 | -5.47 | 0.007 | 3.97 | 0.054 | 3.42 | 0.062 |
| Otu00137 | Bacteria_unclassified | -11.52 | 1.10E-13 | -3.97 | 0.055 | -7.55 | 2.61E-06 | 4.35 | 0.009 |
| Otu00141 | Flavobacterium | -4.54 | 0.025 | -7.26 | 2.89E-04 | 2.73 | 0.144 | 4.40 | 0.015 |
| Otu00142 | Xanthomonadaceae_unclassified | -10.86 | 3.29E-08 | -1.36 | 0.551 | -9.50 | 7.64E-06 | 0.28 | 0.896 |
| Otu00143 | Flavobacterium | -4.15 | 0.022 | -5.36 | 0.003 | 1.21 | 0.470 | 0.93 | 0.587 |
| Otu00144 | Lactobacillus | 5.23 | 1.25E-04 | -2.49 | 0.019 | 7.73 | 4.11E-09 | 2.88 | 0.015 |
| Otu00145 | Bacillus | -7.79 | 0.007 | -0.72 | 0.806 | -7.07 | 0.023 | 4.33 | 0.129 |
| Otu00147 | Aeromonas | 2.98 | 0.104 | -2.04 | 0.281 | 5.02 | 0.005 | -3.74 | 0.019 |
| Otu00150 | Rhodobacter | -6.23 | 3.65E-06 | -2.54 | 0.141 | -3.69 | 0.017 | -1.11 | 0.442 |
| Otu00152 | Leptothrix | -9.75 | 1.36E-08 | -3.77 | 0.086 | -5.98 | 6.33E-04 | 4.24 | 0.015 |
| Otu00154 | Luteolibacter | -11.32 | 8.59E-11 | -4.87 | 0.027 | -6.45 | 1.89E-04 | 0.84 | 0.650 |
| Otu00155 | Rhizobiales_unclassified | -5.60 | 3.38E-04 | -0.11 | 0.947 | -5.50 | 0.002 | 0.49 | 0.782 |
| Otu00156 | Reyranella | 0.18 | 0.871 | -0.65 | 0.484 | 0.84 | 0.394 | 0.74 | 0.442 |
| Otu00157 | Acinetobacter | -0.56 | 0.729 | -2.80 | 0.069 | 2.25 | 0.127 | 2.69 | 0.041 |
| Otu00162 | Rhizobiales_unclassified | -11.44 | 2.55E-12 | -0.89 | 0.659 | -10.54 | 5.75E-09 | -0.11 | 0.958 |
| Otu00163 | Mycobacterium | 2.31 | 0.082 | -2.74 | 0.037 | 5.04 | 4.62E-05 | 0.20 | 0.870 |
| Otu00165 | Brevibacterium | 5.59 | 0.002 | -4.16 | 0.005 | 9.75 | 5.81E-09 | 2.43 | 0.123 |
| Otu00167 | Rhodobacter | -1.10 | 0.537 | -2.97 | 0.095 | 1.86 | 0.248 | 0.15 | 0.926 |
| Otu00168 | Methylobacterium | 1.39 | 0.349 | -0.62 | 0.624 | 2.01 | 0.154 | 3.10 | 0.015 |
| Otu00169 | Arthrobacter | -0.93 | 0.562 | -4.39 | 0.003 | 3.45 | 0.022 | 0.53 | 0.713 |
| Otu00173 | Lactobacillus | 7.01 | 2.97E-06 | -2.24 | 0.072 | 9.25 | 2.12E-10 | 2.64 | 0.042 |
| Otu00176 | Beijerinckiaceae_unclassified | -13.86 | 4.15E-15 | -3.28 | 0.201 | -10.58 | 1.94E-09 | 2.80 | 0.139 |
| Otu00181 | TM7 | -9.41 | 4.87E-08 | -5.18 | 0.011 | -4.24 | 0.013 | 2.42 | 0.163 |
| Otu00182 | Desulfovibrio | 6.77 | 0.003 | -1.79 | 0.407 | 8.56 | 1.25E-04 | -0.72 | 0.743 |
| Otu00183 | Dechloromonas | -8.98 | 3.38E-06 | -8.10 | 1.18E-04 | -0.88 | 0.605 | 0.85 | 0.657 |
| Otu00184 | Leuconostoc | 6.77 | 8.14E-06 | -2.56 | 0.040 | 9.33 | 2.12E-10 | 2.74 | 0.037 |
| Otu00185 | Brevinemataceae_unclassified | 4.89 | 0.032 | -3.14 | 0.185 | 8.03 | 2.30E-04 | 6.48 | 0.001 |
| Otu00192 | Brevinemataceae_unclassified | 8.86 | 1.41E-04 | -3.12 | 0.198 | 11.98 | 1.10E-07 | 4.45 | 0.037 |
| Otu00194 | Psychrobacter | -1.94 | 0.561 | -11.27 | 8.26E-05 | 9.33 | 7.72E-04 | 0.55 | 0.850 |
| Otu00198 | Rhodobacteraceae_unclassified | -7.31 | 5.38E-05 | -1.17 | 0.548 | -6.14 | 0.002 | -0.31 | 0.871 |
| Otu00202 | Bacillus | -6.57 | 6.02E-04 | -4.30 | 0.045 | -2.28 | 0.240 | 0.83 | 0.667 |
| Otu00203 | C111_unclassified | -9.90 | 6.01E-06 | -7.05 | 0.003 | -2.86 | 0.181 | -0.87 | 0.695 |
| Otu00209 | Rhizobiales_unclassified | -7.67 | 9.39E-08 | -5.30 | 0.001 | -2.37 | 0.124 | 0.42 | 0.791 |
| Otu00212 | Rhizobiales_unclassified | -11.16 | 4.26E-09 | -3.30 | 0.213 | -7.86 | 4.14E-05 | 1.50 | 0.464 |

| | | | | | | | | | |
|---|---|---|---|---|---|---|---|---|---|
| Otu00217 | Zymomonas | -6.90 | 4.96E-05 | -1.43 | 0.448 | -5.47 | 0.005 | 0.07 | 0.976 |
| Otu00219 | Mycobacterium | -2.45 | 0.139 | -3.60 | 0.037 | 1.15 | 0.454 | 0.44 | 0.788 |
| Otu00221 | SC | -10.51 | 2.37E-08 | -5.24 | 0.023 | -5.26 | 0.005 | -0.51 | 0.802 |
| Otu00225 | Bacteria_unclassified | -11.31 | 3.11E-04 | -1.68 | 0.605 | -9.63 | 0.004 | 0.95 | 0.782 |
| Otu00226 | Aphanocapsa | -9.04 | 3.29E-09 | 1.11 | 0.564 | -10.15 | 4.43E-08 | 2.88 | 0.098 |
| Otu00227 | Comamonadaceae_unclassified | -10.47 | 1.10E-08 | -3.03 | 0.226 | -7.44 | 9.38E-05 | 1.74 | 0.396 |
| Otu00229 | Staphylococcus | 5.02 | 0.006 | -4.05 | 0.011 | 9.06 | 1.36E-07 | 2.93 | 0.063 |
| Otu00241 | Phyllobacteriaceae_unclassified | -8.90 | 2.56E-06 | -8.40 | 3.48E-05 | -0.49 | 0.762 | -1.70 | 0.383 |
| Otu00247 | Paucibacter | -5.67 | 0.009 | -5.43 | 0.021 | -0.24 | 0.905 | 0.60 | 0.790 |
| Otu00255 | Bacillus | -10.38 | 3.11E-04 | -4.98 | 0.155 | -5.40 | 0.068 | 2.94 | 0.337 |
| Otu00256 | Rhodobacteraceae_unclassified | -12.27 | 1.70E-10 | -5.10 | 0.039 | -7.17 | 8.86E-05 | 0.82 | 0.683 |
| Otu00257 | Brevinemataceae_unclassified | 8.03 | 0.001 | -4.37 | 0.075 | 12.41 | 1.33E-07 | 4.66 | 0.036 |
| Otu00258 | Alphaproteobacteria_unclassified | -4.35 | 0.026 | -9.29 | 1.09E-07 | 4.94 | 0.005 | 0.02 | 0.991 |
| Otu00265 | Reyranella | -4.23 | 0.002 | -4.56 | 0.002 | 0.34 | 0.798 | 0.72 | 0.591 |
| Otu00267 | Rickettsia | -10.51 | 0.016 | -3.30 | 0.448 | -7.21 | 0.107 | -3.06 | 0.468 |
| Otu00270 | OM60_unclassified | -11.17 | 1.24E-06 | -3.32 | 0.277 | -7.85 | 9.70E-04 | 1.73 | 0.468 |
| Otu00288 | Hyphomicrobium | -8.77 | 1.01E-06 | -1.94 | 0.401 | -6.83 | 4.38E-04 | -0.36 | 0.854 |
| Otu00291 | Xanthobacteraceae_unclassified | 0.82 | 0.712 | -6.83 | 1.94E-04 | 7.65 | 5.07E-05 | -0.70 | 0.713 |
| Otu00305 | Aeromonadaceae_unclassified | -1.66 | 0.558 | -11.93 | 9.38E-08 | 10.27 | 3.04E-06 | 0.22 | 0.926 |
| Otu00306 | Fusobacteriales_unclassified | -2.80 | 0.349 | -9.79 | 1.90E-04 | 6.99 | 0.006 | -1.64 | 0.503 |
| Otu00321 | BD1 | -5.26 | 0.066 | -8.37 | 0.003 | 3.11 | 0.231 | 0.94 | 0.727 |
| Otu00326 | Planctomyces | -8.16 | 1.64E-06 | -4.36 | 0.039 | -3.80 | 0.043 | -0.28 | 0.871 |
| Otu00331 | Bacteria_unclassified | -11.94 | 3.17E-11 | -3.28 | 0.201 | -8.65 | 1.67E-06 | 2.38 | 0.234 |
| Otu00332 | Fusibacter | -2.91 | 0.407 | -9.65 | 0.002 | 6.74 | 0.028 | -1.21 | 0.688 |
| Otu00338 | Chthoniobacter | -6.06 | 0.002 | -0.57 | 0.788 | -5.49 | 0.014 | 1.17 | 0.570 |
| Otu00340 | Rhizobiales_unclassified | -4.70 | 0.013 | -7.14 | 1.67E-04 | 2.44 | 0.173 | 2.07 | 0.258 |
| Otu00342 | Betaproteobacteria_unclassified | -8.95 | 0.006 | -3.30 | 0.379 | -5.65 | 0.093 | 3.70 | 0.265 |
| Otu00351 | Planctomyces | -10.69 | 9.25E-08 | -5.58 | 0.023 | -5.11 | 0.009 | 0.50 | 0.819 |
| Otu00354 | Rhodobacter | -11.02 | 1.34E-10 | -1.96 | 0.405 | -9.05 | 1.06E-06 | 3.37 | 0.064 |
| Otu00367 | Gemmata | -11.29 | 2.86E-09 | -3.29 | 0.213 | -8.00 | 2.87E-05 | 2.45 | 0.240 |
| Otu00374 | Gemmataceae_unclassified | -9.24 | 8.53E-07 | -5.07 | 0.028 | -4.17 | 0.028 | 1.00 | 0.601 |
| Otu00393 | Luteolibacter | -11.17 | 6.69E-08 | -5.22 | 0.043 | -5.95 | 0.004 | 0.44 | 0.843 |
| Otu00399 | Roseococcus | -8.66 | 3.12E-07 | -1.68 | 0.447 | -6.99 | 2.03E-04 | 3.62 | 0.044 |
| Otu00425 | Moraxellaceae_unclassified | -8.86 | 0.009 | -3.26 | 0.397 | -5.59 | 0.108 | 4.82 | 0.139 |
| Otu00428 | Gemmata | -7.32 | 1.04E-06 | -4.57 | 0.009 | -2.75 | 0.093 | 0.22 | 0.889 |
| Otu00445 | Methylobacteriaceae_unclassified | -12.09 | 3.05E-11 | -4.75 | 0.046 | -7.34 | 1.86E-05 | 0.80 | 0.679 |
| Otu00454 | Clostridium | -5.37 | 0.091 | -9.26 | 0.003 | 3.89 | 0.173 | -0.33 | 0.909 |
| Otu00470 | DA101 | -8.20 | 6.33E-05 | -2.82 | 0.289 | -5.37 | 0.015 | 2.63 | 0.234 |
| Otu00482 | Brevinemataceae_unclassified | 3.80 | 0.141 | -2.94 | 0.268 | 6.74 | 0.007 | 5.85 | 0.011 |

| | | | | | | | | | |
|---|---|---|---|---|---|---|---|---|---|
| Otu00484 | Gemmata | -9.54 | 1.00E-06 | -2.41 | 0.358 | -7.14 | 6.65E-04 | 3.21 | 0.117 |
| Otu00496 | Turicibacter | -9.53 | 6.69E-06 | -7.71 | 8.24E-04 | -1.82 | 0.326 | 1.45 | 0.470 |
| Otu00532 | Gammaproteobacteria_unclassified | -10.79 | 1.27E-04 | -3.32 | 0.343 | -7.47 | 0.011 | 2.56 | 0.396 |
| Otu00543 | Bacteria_unclassified | -10.91 | 2.22E-08 | -3.29 | 0.221 | -7.61 | 1.23E-04 | 2.68 | 0.202 |
| Otu00546 | Brevinemataceae_unclassified | 2.19 | 0.428 | -3.41 | 0.213 | 5.59 | 0.030 | 5.10 | 0.027 |
| Otu00552 | Brevinemataceae_unclassified | 2.82 | 0.306 | -3.74 | 0.177 | 6.55 | 0.012 | 6.53 | 0.006 |
| Otu00581 | Bacteria_unclassified | -10.32 | 1.04E-05 | -3.31 | 0.281 | -7.01 | 0.004 | 2.68 | 0.293 |
| Otu00585 | Bacteria_unclassified | -11.15 | 4.26E-09 | -3.29 | 0.213 | -7.86 | 4.25E-05 | 2.23 | 0.293 |
| Otu00586 | Bacteria_unclassified | -10.73 | 3.71E-08 | -4.19 | 0.105 | -6.53 | 7.26E-04 | 3.01 | 0.129 |
| Otu00588 | Pirellulaceae_unclassified | -10.59 | 1.68E-06 | -6.46 | 0.012 | -4.14 | 0.063 | -0.04 | 0.989 |
| Otu00630 | Bacteria_unclassified | -10.80 | 1.04E-07 | -3.30 | 0.234 | -7.50 | 3.27E-04 | 2.45 | 0.271 |
| Otu00667 | Brevinemataceae_unclassified | 6.45 | 0.008 | -4.00 | 0.094 | 10.45 | 6.26E-06 | 4.35 | 0.043 |
| Otu00668 | Brevinemataceae_unclassified | 3.52 | 0.173 | -3.23 | 0.213 | 6.75 | 0.006 | 5.49 | 0.015 |
| Otu00678 | Bacillales_unclassified | -2.79 | 0.412 | -12.53 | 1.38E-05 | 9.74 | 4.19E-04 | 1.33 | 0.640 |
| Otu00691 | Bacteria_unclassified | -9.71 | 1.41E-06 | -4.20 | 0.112 | -5.51 | 0.007 | 3.23 | 0.110 |
| Otu00734 | Ruminococcaceae_unclassified | -11.88 | 0.001 | -3.39 | 0.408 | -8.49 | 0.024 | -0.26 | 0.953 |
| Otu00748 | Roseomonas | -9.71 | 5.33E-07 | -2.75 | 0.298 | -6.96 | 6.72E-04 | 3.53 | 0.071 |
| Otu00758 | Bacteria_unclassified | -10.47 | 9.39E-08 | -3.30 | 0.223 | -7.18 | 3.64E-04 | 2.35 | 0.279 |
| Otu00780 | Brevinemataceae_unclassified | 3.60 | 0.158 | -2.91 | 0.262 | 6.51 | 0.008 | 5.32 | 0.017 |
| Otu00821 | Brevinemataceae_unclassified | 3.36 | 0.180 | -3.14 | 0.209 | 6.50 | 0.006 | 5.71 | 0.010 |
| Otu00826 | Gemmata | -11.28 | 7.42E-07 | -3.32 | 0.274 | -7.96 | 6.81E-04 | 1.92 | 0.419 |
| Otu00917 | Ruminococcaceae_unclassified | -11.55 | 1.60E-04 | -3.38 | 0.363 | -8.17 | 0.011 | -0.09 | 0.979 |
| Otu00942 | Gemmata | -8.74 | 6.34E-06 | -2.86 | 0.264 | -5.88 | 0.005 | 2.75 | 0.182 |
| Otu00956 | Brevinemataceae_unclassified | 2.71 | 0.349 | -3.83 | 0.181 | 6.54 | 0.017 | 6.35 | 0.011 |
| Otu00978 | SHA | -2.75 | 0.429 | -10.97 | 2.62E-04 | 8.22 | 0.005 | 2.22 | 0.442 |
| Otu00991 | Rhizobiales_unclassified | -10.30 | 7.29E-08 | -4.19 | 0.099 | -6.10 | 0.001 | 1.64 | 0.413 |
| Otu00994 | Bacteria_unclassified | -11.19 | 3.16E-06 | -3.32 | 0.291 | -7.87 | 0.002 | 2.17 | 0.396 |
| Otu01053 | Brevinemataceae_unclassified | 2.21 | 0.454 | -3.74 | 0.193 | 5.95 | 0.030 | 6.45 | 0.010 |
| Otu01109 | Brevinemataceae_unclassified | 2.08 | 0.473 | -4.01 | 0.155 | 6.10 | 0.023 | 6.63 | 0.006 |
| Otu01177 | Bacteria_unclassified | -7.29 | 0.015 | -3.29 | 0.346 | -4.00 | 0.183 | 4.61 | 0.106 |
| Otu01217 | Gammaproteobacteria_unclassified | -9.67 | 0.014 | -3.29 | 0.435 | -6.38 | 0.112 | 3.15 | 0.415 |
| Otu01228 | Rhizobiales_unclassified | -8.62 | 0.004 | -3.31 | 0.348 | -5.31 | 0.087 | 3.31 | 0.281 |
| Otu01232 | Brevinemataceae_unclassified | 3.03 | 0.264 | -2.40 | 0.373 | 5.43 | 0.043 | 5.66 | 0.017 |
| Otu01269 | Enterobacteriaceae_unclassified | -8.93 | 0.025 | -3.29 | 0.434 | -5.64 | 0.156 | 3.13 | 0.416 |
| Otu01393 | Gammaproteobacteria_unclassified | -9.51 | 2.72E-04 | -3.31 | 0.313 | -6.19 | 0.024 | 2.81 | 0.317 |
| Otu01425 | Brevinemataceae_unclassified | 2.55 | 0.329 | -3.40 | 0.183 | 5.95 | 0.016 | 5.34 | 0.016 |
| Otu01461 | Brevinemataceae_unclassified | 2.97 | 0.243 | -2.78 | 0.282 | 5.75 | 0.022 | 5.30 | 0.017 |
| Otu01500 | Gammaproteobacteria_unclassified | -7.99 | 0.076 | -3.22 | 0.448 | -4.77 | 0.269 | 4.15 | 0.364 |
| Otu01686 | Bacteria_unclassified | -8.31 | 0.009 | -3.30 | 0.372 | -5.01 | 0.124 | 3.67 | 0.258 |

| | | | | | | | | | |
|---|---|---|---|---|---|---|---|---|---|
| Otu01794 | Brevinemataceae_unclassified | 1.84 | 0.520 | -4.13 | 0.130 | 5.97 | 0.023 | 4.97 | 0.034 |
| Otu01890 | Gammaproteobacteria_unclassified | -8.24 | 0.066 | -3.22 | 0.448 | -5.02 | 0.246 | 4.08 | 0.375 |
| Otu01981 | Brevinemataceae_unclassified | 1.43 | 0.641 | -4.11 | 0.177 | 5.54 | 0.056 | 5.59 | 0.030 |
| Otu02038 | Gammaproteobacteria_unclassified | -8.75 | 0.028 | -3.28 | 0.434 | -5.46 | 0.166 | 3.28 | 0.402 |
| Otu02218 | Brevinemataceae_unclassified | 2.09 | 0.518 | -3.43 | 0.270 | 5.52 | 0.065 | 5.50 | 0.037 |
| Otu02266 | Brevinemataceae_unclassified | 1.50 | 0.571 | -3.45 | 0.183 | 4.94 | 0.049 | 5.31 | 0.017 |
| Otu02309 | Enterobacteriaceae_unclassified | -9.50 | 0.005 | -4.03 | 0.307 | -5.47 | 0.110 | 2.44 | 0.463 |
| Otu02323 | Brevinemataceae_unclassified | 1.37 | 0.650 | -3.55 | 0.234 | 4.92 | 0.087 | 5.95 | 0.019 |
| Otu02372 | Brevinemataceae_unclassified | 2.04 | 0.510 | -3.72 | 0.207 | 5.76 | 0.044 | 5.24 | 0.037 |
| Otu02448 | Brevinemataceae_unclassified | 1.31 | 0.667 | -4.02 | 0.177 | 5.33 | 0.061 | 6.36 | 0.014 |
| Otu02472 | Brevinemataceae_unclassified | 0.87 | 0.761 | -4.12 | 0.120 | 5.00 | 0.052 | 5.32 | 0.019 |
| Otu02475 | Gammaproteobacteria_unclassified | -9.50 | 0.008 | -3.30 | 0.409 | -6.20 | 0.094 | 3.05 | 0.397 |
| Otu02629 | Brevinemataceae_unclassified | 0.88 | 0.724 | -3.76 | 0.110 | 4.65 | 0.045 | 5.85 | 0.005 |
| Otu02744 | Enterobacteriaceae_unclassified | -9.16 | 0.005 | -3.31 | 0.384 | -5.85 | 0.087 | 3.11 | 0.377 |
| Otu02750 | Brevinemataceae_unclassified | 3.14 | 0.329 | -2.05 | 0.448 | 5.20 | 0.093 | 3.72 | 0.196 |
| Otu02776 | Enterobacteriaceae_unclassified | -7.98 | 0.076 | -3.22 | 0.448 | -4.77 | 0.269 | 4.15 | 0.364 |
| Otu02839 | Enterobacteriaceae_unclassified | -7.90 | 0.068 | -3.24 | 0.448 | -4.66 | 0.264 | 4.00 | 0.364 |
| Otu02936 | Brevinemataceae_unclassified | 1.28 | 0.690 | -4.33 | 0.164 | 5.61 | 0.059 | 5.61 | 0.034 |
| Otu02943 | Brevinemataceae_unclassified | 1.73 | 0.571 | -4.86 | 0.096 | 6.60 | 0.022 | 4.96 | 0.051 |
| Otu02970 | Brevinemataceae_unclassified | 1.55 | 0.635 | -2.83 | 0.373 | 4.37 | 0.154 | 6.40 | 0.019 |
| Otu03195 | Brevinemataceae_unclassified | 1.07 | 0.756 | -4.06 | 0.209 | 5.12 | 0.097 | 5.40 | 0.048 |
| Otu03299 | Brevinemataceae_unclassified | 0.56 | 0.876 | -5.38 | 0.105 | 5.94 | 0.063 | 5.71 | 0.044 |
| Otu03416 | Enterobacteriaceae_unclassified | -8.11 | 0.071 | -3.21 | 0.448 | -4.90 | 0.256 | 4.19 | 0.362 |
| Otu03542 | Brevinemataceae_unclassified | 2.43 | 0.430 | -3.53 | 0.231 | 5.95 | 0.041 | 4.52 | 0.071 |
| Otu04096 | Gammaproteobacteria_unclassified | -6.85 | 0.117 | -2.03 | 0.619 | -4.83 | 0.254 | 3.41 | 0.413 |
| Otu04193 | Brevinemataceae_unclassified | 0.26 | 0.945 | -5.02 | 0.167 | 5.28 | 0.121 | 5.16 | 0.095 |
| Otu04634 | Brevinemataceae_unclassified | -0.77 | 0.847 | -6.15 | 0.095 | 5.37 | 0.117 | 6.44 | 0.039 |

* Fold changes are coloured coded as follows: green= increased abundance in natural treatment, red= increased abundance in hatchery treatment, blue=increased abundance in enriched treatment

**Table S3. Differentially abundant skin OTUs**

| OTU | Annotation | H:W Log2 Fold Change | H:W FDR | H:E Log2 Fold Change | H:E FDR | E:W Log2 Fold Change | E:W FDR | H:W origin Log2 Fold Change | Origin FDR |
|---|---|---|---|---|---|---|---|---|---|
| Otu00002 | Alphaproteobacteria_unclassified | 1.556376 | 0.044 | -0.6488 | 0.952 | 2.205177 | 0.020 | 2.778817 | 0.001 |
| Otu00008 | Achromobacter | 0.773597 | 0.557 | -1.86711 | 0.451 | 2.64071 | 0.024 | -1.39934 | 0.661 |
| Otu00010 | Aeromonadaceae_unclassified | -0.83847 | 0.609 | -4.33035 | 0.007 | 3.491887 | 0.015 | -0.93435 | 0.912 |
| Otu00014 | Staphylococcus | -0.80568 | 0.132 | 0.552315 | 0.904 | -1.358 | 0.015 | -0.39514 | 0.912 |
| Otu00015 | Enterobacteriaceae_unclassified | 4.188617 | 1.05E-05 | 0.010241 | 0.994 | 4.178375 | 1.73E-05 | -1.57605 | 0.351 |
| Otu00018 | Rhodoferax | -5.56918 | 3.04E-10 | 0.389738 | 0.952 | -5.95891 | 8.10E-12 | -0.84684 | 0.887 |
| Otu00021 | Clostridium | 2.386047 | 0.047 | 0.735431 | 0.952 | 1.650615 | 0.201 | 0.661986 | 0.919 |
| Otu00024 | Pseudomonas | 1.709872 | 0.045 | -0.15634 | 0.952 | 1.866213 | 0.040 | 0.070525 | 0.999 |
| Otu00026 | Deefgea | -1.46914 | 0.553 | -5.79485 | 0.022 | 4.325711 | 0.029 | 1.387377 | 0.912 |
| Otu00030 | Burkholderia | 5.202669 | 0.033 | -4.33959 | 0.287 | 9.542258 | 4.84E-05 | -3.6526 | 0.351 |
| Otu00031 | Neisseriaceae_unclassified | 10.28094 | 1.05E-05 | 10.57511 | 1.14E-05 | -0.29417 | 0.935 | -1.38232 | 0.912 |
| Otu00036 | Rhodococcus | -2.36846 | 0.003 | 0.802573 | 0.939 | -3.17103 | 4.56E-05 | 0.011785 | 0.999 |
| Otu00039 | Rhizobium | -3.31869 | 0.119 | 1.610136 | 0.952 | -4.92882 | 0.033 | -1.77728 | 0.912 |
| Otu00041 | Alphaproteobacteria_unclassified | 9.182587 | 1.27E-08 | -1.94119 | 0.645 | 11.12378 | 1.58E-12 | -3.03152 | 0.194 |
| Otu00046 | Enterobacter | 2.525711 | 0.033 | -0.44464 | 0.952 | 2.970353 | 0.015 | -0.041 | 0.999 |
| Otu00052 | Sphingomonas | -4.2006 | 3.22E-04 | -0.46794 | 0.952 | -3.73266 | 0.002 | -0.32245 | 0.999 |
| Otu00053 | Rhodococcus | -2.31309 | 0.003 | -0.39538 | 0.952 | -1.91771 | 0.019 | -0.78072 | 0.777 |
| Otu00056 | Comamonadaceae_unclassified | -4.04317 | 4.14E-07 | -0.47741 | 0.952 | -3.56576 | 1.73E-05 | -0.70796 | 0.912 |
| Otu00060 | Variovorax | -2.11098 | 0.004 | 0.565638 | 0.952 | -2.67662 | 2.11E-04 | -0.41798 | 0.912 |
| Otu00068 | Rhodobacter | -4.54138 | 0.014 | -0.85512 | 0.952 | -3.68626 | 0.054 | -1.60721 | 0.887 |
| Otu00069 | Rhodoferax | -2.04351 | 0.272 | 3.065127 | 0.417 | -5.10864 | 0.005 | -0.37976 | 0.999 |
| Otu00077 | Sphingomonas | -5.24101 | 1.20E-04 | -0.84465 | 0.952 | -4.39636 | 0.002 | 0.140824 | 0.999 |
| Otu00078 | Pseudomonas | -0.68157 | 0.602 | -2.99821 | 0.030 | 2.31664 | 0.045 | 0.348338 | 0.999 |
| Otu00083 | Aeromonadaceae_unclassified | -2.48151 | 0.335 | -8.40767 | 4.85E-04 | 5.926166 | 0.011 | -1.55145 | 0.912 |
| Otu00085 | Limnohabitans | -13.8922 | 3.04E-10 | -0.75123 | 0.952 | -13.141 | 1.96E-09 | -0.16489 | 0.999 |
| Otu00086 | Betaproteobacteria_unclassified | -6.33765 | 0.013 | -3.95385 | 0.480 | -2.3838 | 0.361 | 2.985286 | 0.661 |
| Otu00093 | Bosea | -11.3897 | 1.05E-05 | -0.75169 | 0.952 | -10.638 | 4.56E-05 | 0.10814 | 0.999 |
| Otu00102 | Agrobacterium | -5.59641 | 3.33E-04 | -2.05423 | 0.716 | -3.54218 | 0.034 | -1.40794 | 0.887 |
| Otu00109 | Flavobacterium | -6.16318 | 0.001 | -7.7421 | 1.21E-04 | 1.57892 | 0.402 | -3.34525 | 0.194 |
| Otu00112 | Sinobacteraceae_unclassified | -3.99249 | 0.003 | 3.624325 | 0.146 | -7.61681 | 8.61E-07 | -0.56716 | 0.999 |
| Otu00115 | Proteobacteria_unclassified | 10.38305 | 2.51E-06 | 7.558698 | 4.85E-04 | 2.824348 | 0.289 | -0.95407 | 0.999 |
| Otu00118 | Oxalobacteraceae_unclassified | -4.32887 | 0.003 | 0.44615 | 0.952 | -4.77502 | 0.001 | -1.06688 | 0.912 |
| Otu00120 | Procabacteriaceae_unclassified | 7.733133 | 0.001 | -2.44839 | 0.777 | 10.18153 | 1.73E-05 | 2.513067 | 0.683 |
| Otu00123 | Pelomonas | -2.68044 | 0.040 | 2.212309 | 0.464 | -4.89275 | 1.93E-04 | -1.18499 | 0.887 |
| Otu00127 | Caulobacteraceae_unclassified | -2.71234 | 0.033 | -0.97336 | 0.952 | -1.73898 | 0.199 | -0.17175 | 0.999 |

| | | | | | | | | | |
|---|---|---|---|---|---|---|---|---|---|
| Otu00129 | Flectobacillus | -5.76928 | 0.005 | 0.93251 | 0.952 | -6.70179 | 0.003 | -4.60311 | 0.067 |
| Otu00130 | Comamonadaceae_unclassified | -4.19369 | 0.001 | -1.52503 | 0.777 | -2.66866 | 0.049 | -0.03595 | 0.999 |
| Otu00131 | Novosphingobium | -4.76893 | 0.020 | -0.89846 | 0.952 | -3.87047 | 0.067 | -2.04148 | 0.777 |
| Otu00133 | Brevundimonas | -3.84314 | 2.53E-04 | -0.82492 | 0.952 | -3.01823 | 0.006 | -0.66597 | 0.912 |
| Otu00138 | Betaproteobacteria_unclassified | -5.13204 | 0.001 | 1.459914 | 0.952 | -6.59196 | 2.14E-05 | -1.53903 | 0.887 |
| Otu00141 | Flavobacterium | -9.40745 | 7.75E-06 | -10.3137 | 2.20E-06 | 0.906216 | 0.680 | -1.32151 | 0.912 |
| Otu00146 | Rudanella | 4.175492 | 0.020 | 1.043143 | 0.952 | 3.132349 | 0.088 | -2.12046 | 0.661 |
| Otu00147 | Aeromonas | -3.22424 | 0.047 | -4.34954 | 0.038 | 1.125297 | 0.566 | -1.12602 | 0.912 |
| Otu00148 | Enterobacteriaceae_unclassified | 5.162619 | 0.001 | -1.09014 | 0.952 | 6.25276 | 8.54E-05 | -2.15939 | 0.591 |
| Otu00152 | Leptothrix | -6.64104 | 0.001 | -5.09079 | 0.071 | -1.55025 | 0.469 | -2.63125 | 0.591 |
| Otu00161 | Polaromonas | -6.60331 | 0.001 | -3.59836 | 0.396 | -3.00495 | 0.112 | -2.3595 | 0.661 |
| Otu00165 | Brevibacterium | -6.04605 | 0.031 | -10.1922 | 4.85E-04 | 4.146168 | 0.094 | 0.208692 | 0.999 |
| Otu00183 | Dechloromonas | -5.07152 | 0.013 | -6.84333 | 0.002 | 1.771806 | 0.376 | 0.609082 | 0.999 |
| Otu00199 | Oxalobacteraceae_unclassified | -4.15169 | 0.069 | -11.7713 | 3.91E-09 | 7.619628 | 1.97E-05 | -2.56207 | 0.661 |
| Otu00201 | Methylibium | -3.37965 | 0.096 | 0.962008 | 0.952 | -4.34166 | 0.045 | -0.61099 | 0.999 |
| Otu00212 | Rhizobiales_unclassified | -9.49564 | 3.23E-04 | 0.555565 | 0.952 | -10.0512 | 2.11E-04 | -1.70166 | 0.912 |
| Otu00238 | Aquabacterium | -2.61753 | 0.220 | 1.999647 | 0.952 | -4.61718 | 0.037 | -0.31329 | 0.999 |
| Otu00247 | Paucibacter | -5.81289 | 0.013 | -6.34086 | 0.029 | 0.527976 | 0.863 | -2.00785 | 0.887 |
| Otu00292 | Rhodoferax | -7.00651 | 0.007 | -3.06174 | 0.777 | -3.94477 | 0.120 | 1.255962 | 0.957 |
| Otu00294 | Chryseobacterium | -5.06418 | 0.033 | -6.81045 | 0.021 | 1.746277 | 0.497 | 1.500284 | 0.912 |
| Otu00299 | Ellin6075_unclassified | -8.11813 | 0.002 | -3.31148 | 0.758 | -4.80665 | 0.067 | -1.94716 | 0.912 |
| Otu00309 | Oxalobacteraceae_unclassified | 3.272396 | 0.213 | -3.11094 | 0.683 | 6.383337 | 0.016 | -2.6458 | 0.707 |
| Otu00317 | Sphingomonadaceae_unclassified | -10.5568 | 0.003 | 0.395469 | 0.966 | -10.9523 | 0.002 | -0.74326 | 0.999 |
| Otu00323 | Leadbetterella | -7.56495 | 1.07E-04 | -0.91272 | 0.952 | -6.65223 | 0.001 | -1.30059 | 0.912 |
| Otu00341 | Cyanobacteria_unclassified | -15.0992 | 0.001 | -10.0454 | 0.146 | -5.05385 | 0.286 | -1.01737 | 0.999 |
| Otu00342 | Betaproteobacteria_unclassified | -10.6483 | 3.23E-05 | -0.74739 | 0.952 | -9.90087 | 1.41E-04 | -0.77514 | 0.999 |
| Otu00359 | Sediminibacterium | -5.05367 | 0.022 | -4.3068 | 0.257 | -0.74686 | 0.780 | -3.76851 | 0.197 |
| Otu00368 | Spirosoma | -10.8473 | 2.17E-07 | -0.75168 | 0.952 | -10.0956 | 2.12E-06 | -0.11562 | 0.999 |
| Otu00372 | Alkanindiges | -8.41141 | 8.11E-05 | 0.15676 | 0.984 | -8.56817 | 9.25E-05 | 0.397859 | 0.999 |
| Otu00400 | Collimonas | -5.64549 | 0.016 | -2.20158 | 0.952 | -3.44391 | 0.138 | 0.757283 | 0.999 |
| Otu00410 | Chamaesiphonaceae_unclassified | -10.7357 | 7.91E-06 | -0.75136 | 0.952 | -9.98436 | 4.13E-05 | -0.1294 | 0.999 |
| Otu00422 | Chitinophagaceae_unclassified | -10.9877 | 0.011 | -0.73685 | 0.952 | -10.2508 | 0.021 | -1.06434 | 0.999 |
| Otu00432 | Pirellulaceae_unclassified | -9.82841 | 1.16E-05 | 0.277167 | 0.966 | -10.1056 | 1.82E-05 | 0.561771 | 0.999 |
| Otu00434 | Dyadobacter | -11.1624 | 4.90E-07 | -3.54565 | 0.606 | -7.61677 | 2.94E-04 | -2.33261 | 0.777 |
| Otu00436 | Pedobacter | -7.09587 | 0.019 | -1.56624 | 0.952 | -5.52962 | 0.067 | 0.703913 | 0.999 |
| Otu00469 | Spirosoma | -11.7358 | 0.001 | -0.75052 | 0.952 | -10.9853 | 0.002 | -0.00362 | 0.999 |
| Otu00489 | Betaproteobacteria_unclassified | -10.5689 | 1.44E-05 | -0.75057 | 0.952 | -9.8183 | 7.86E-05 | -0.27071 | 0.999 |
| Otu00502 | Solimonas | -10.9978 | 3.23E-04 | -0.74979 | 0.952 | -10.248 | 0.001 | -0.26394 | 0.999 |
| Otu00523 | Methylotenera | -10.5077 | 0.003 | -0.73708 | 0.952 | -9.77064 | 0.008 | -1.39182 | 0.999 |

| | | | | | | | | | |
|---|---|---|---|---|---|---|---|---|---|
| Otu00524 | Saprospirales | -8.47408 | 0.035 | -0.73013 | 0.952 | -7.74394 | 0.067 | -2.90224 | 0.912 |
| Otu00558 | Rudanella | -11.4514 | 0.004 | -0.74261 | 0.952 | -10.7088 | 0.010 | -0.73085 | 0.999 |
| Otu00609 | Cytophagales_unclassified | -8.60899 | 0.006 | 0.830282 | 0.952 | -9.43927 | 0.004 | -1.98804 | 0.912 |
| Otu00612 | Comamonadaceae_unclassified | -9.70239 | 0.031 | -4.74063 | 0.881 | -4.96177 | 0.285 | 0.403652 | 0.999 |
| Otu00621 | Polyangiaceae_unclassified | -8.2212 | 0.002 | -1.27302 | 0.952 | -6.94818 | 0.012 | -4.26754 | 0.351 |
| Otu00629 | Devosia | -7.06167 | 0.049 | -3.01464 | 0.952 | -4.04702 | 0.291 | 0.879318 | 0.999 |
| Otu00681 | Pedobacter | -7.08363 | 0.033 | 1.92324 | 0.952 | -9.00687 | 0.013 | 0.18831 | 0.999 |
| Otu00685 | Nevskia | -6.8355 | 0.050 | 2.87173 | 0.952 | -9.70723 | 0.011 | -3.63723 | 0.777 |
| Otu00699 | Betaproteobacteria_unclassified | -10.1741 | 0.004 | -5.40188 | 0.505 | -4.7722 | 0.160 | 0.666321 | 0.999 |
| Otu01041 | Spirosoma | -9.82048 | 0.021 | -0.73331 | 0.952 | -9.08717 | 0.041 | -1.35519 | 0.999 |

* Fold changes are coloured coded as follows: green= increased abundance in natural treatment, red= increased abundance in hatchery treatment, blue=increased abundance in enriched treatment

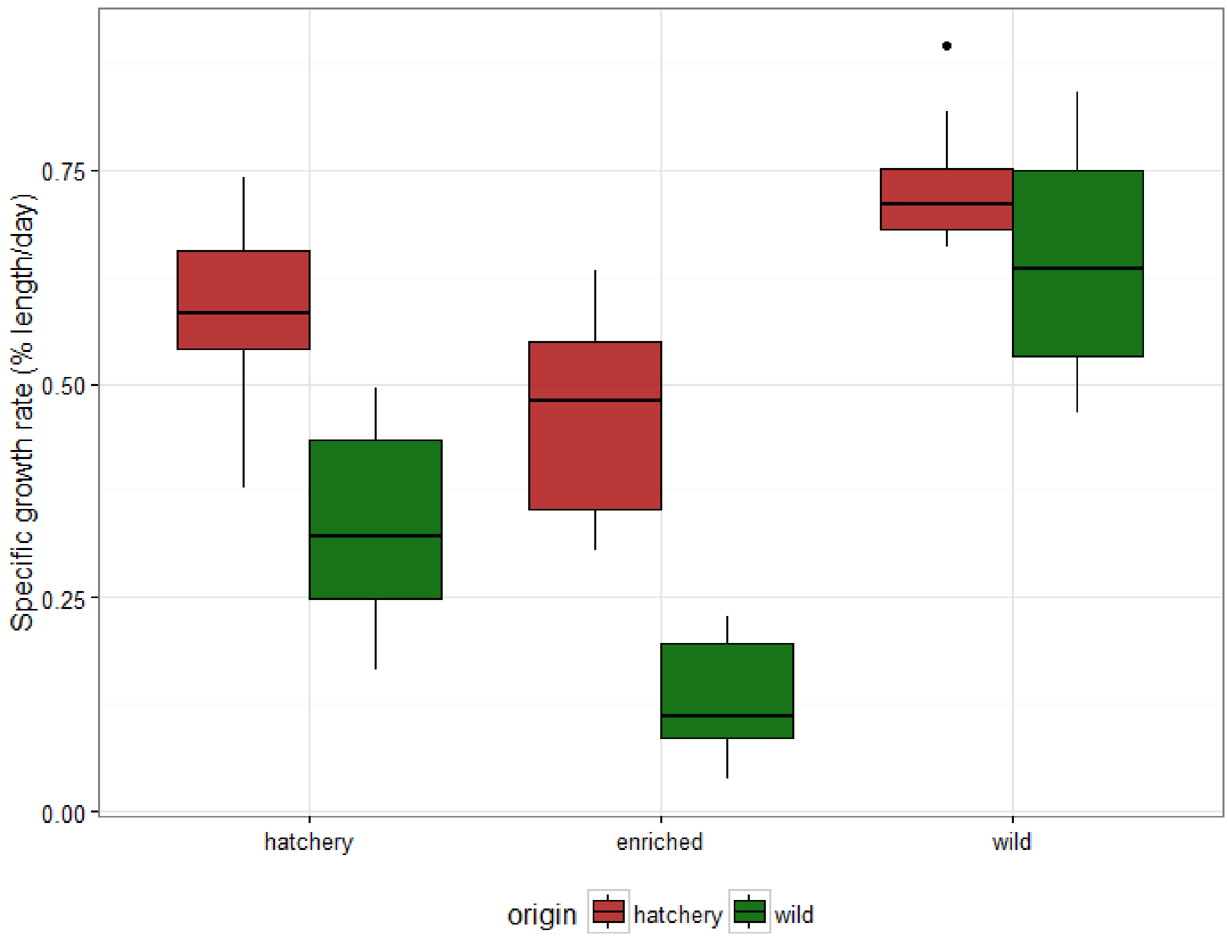


**Figure S1.** Specific growth rate for individually matched fish during the course of the experiment of fish in each treatment group (n=8).

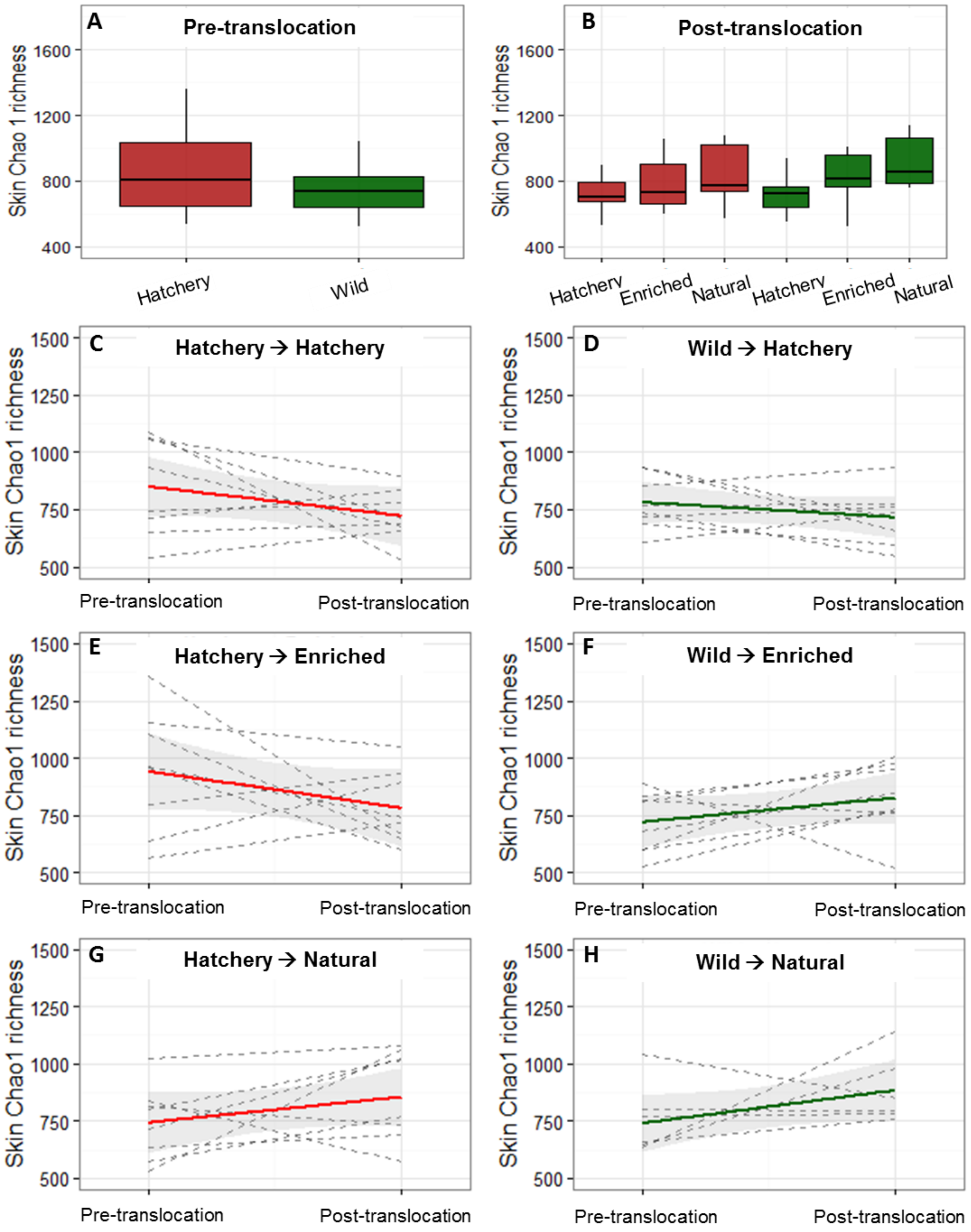


**Figure S2.** Skin Chao1 richness in each group A) before (n=24) and B) after the experiment (n=8), red shading indicates hatchery origin and green shading indicates wild origin fish. C-H) Change in faecal Chao1 richness for all matched fish during the course of the experiment, each dashed line represents an individual fish, with coloured lines displaying the average for each group and grey shading indicating 95% confidence intervals.

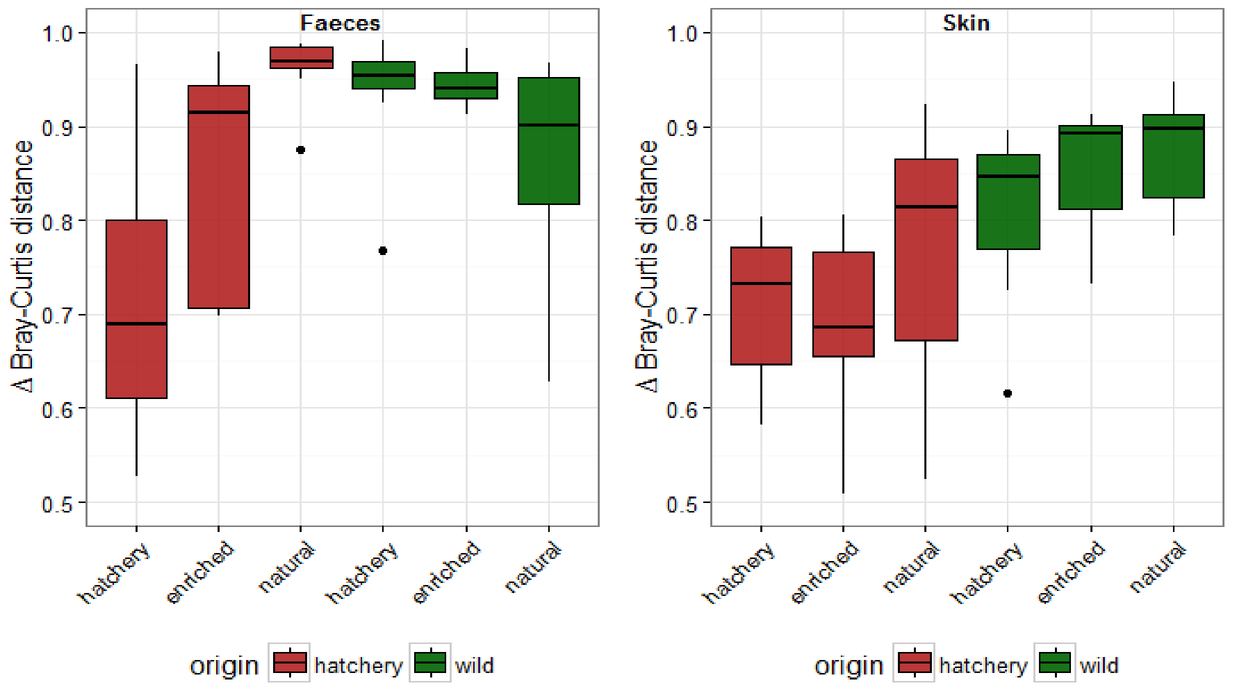


**Figure S3.** Change in microbiome structure for individually matched fish over the course of the experiment, based on Bray-Curtis distances, in each treatment group (n=8). Δ Bray-Curtis distance= Bray-Curtis dissimilarity index post-translocation (ii) - Bray-Curtis dissimilarity index pre-translocation (i).

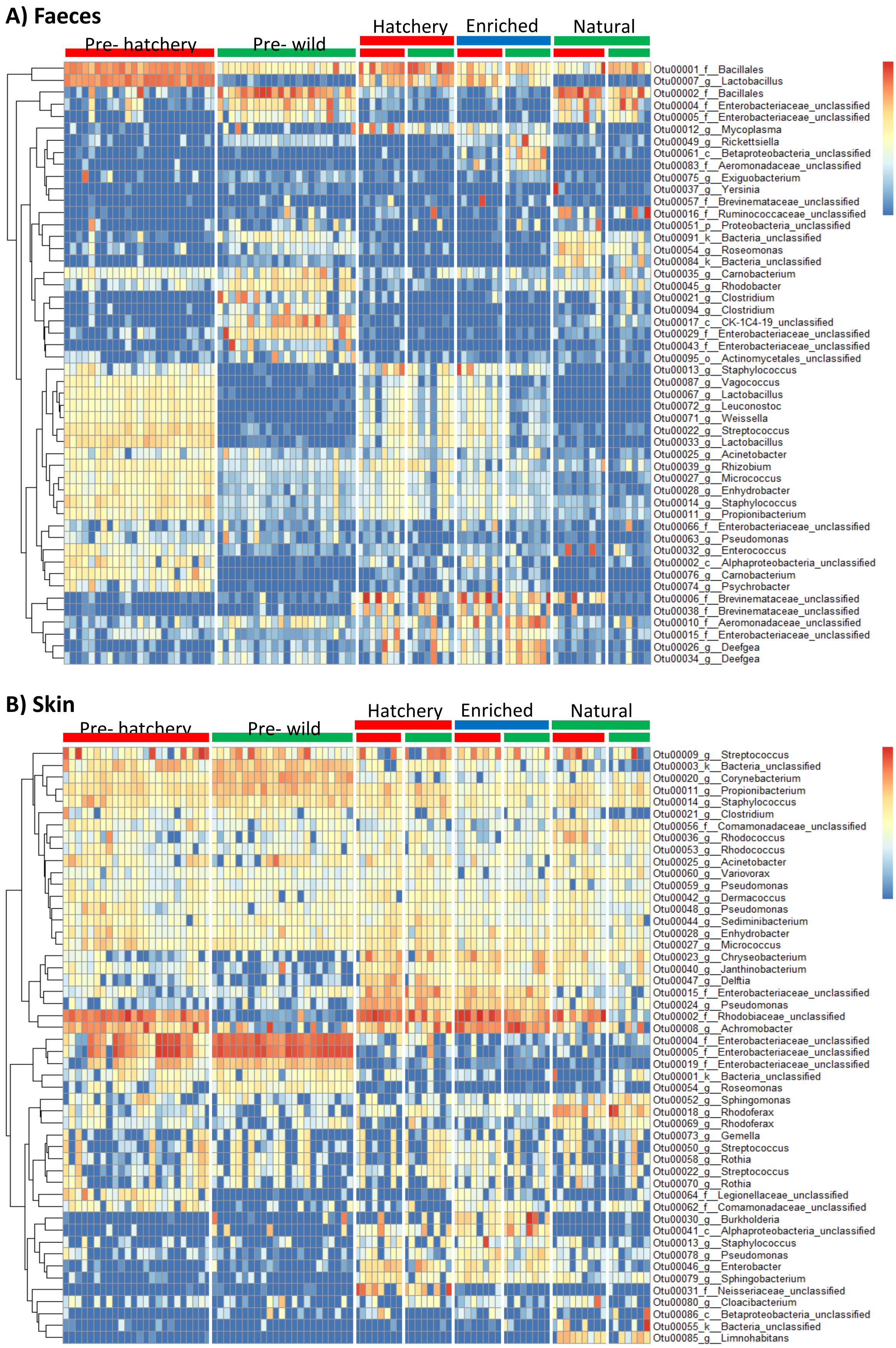


**Figure S4.** Heatmap illustrating the abundance of the top 50 OTUs in the gut and in the skin microbiome. Data presented are log2 transformed OTU read counts, and hierarchical clustering was based on an Euclidean distance metric.

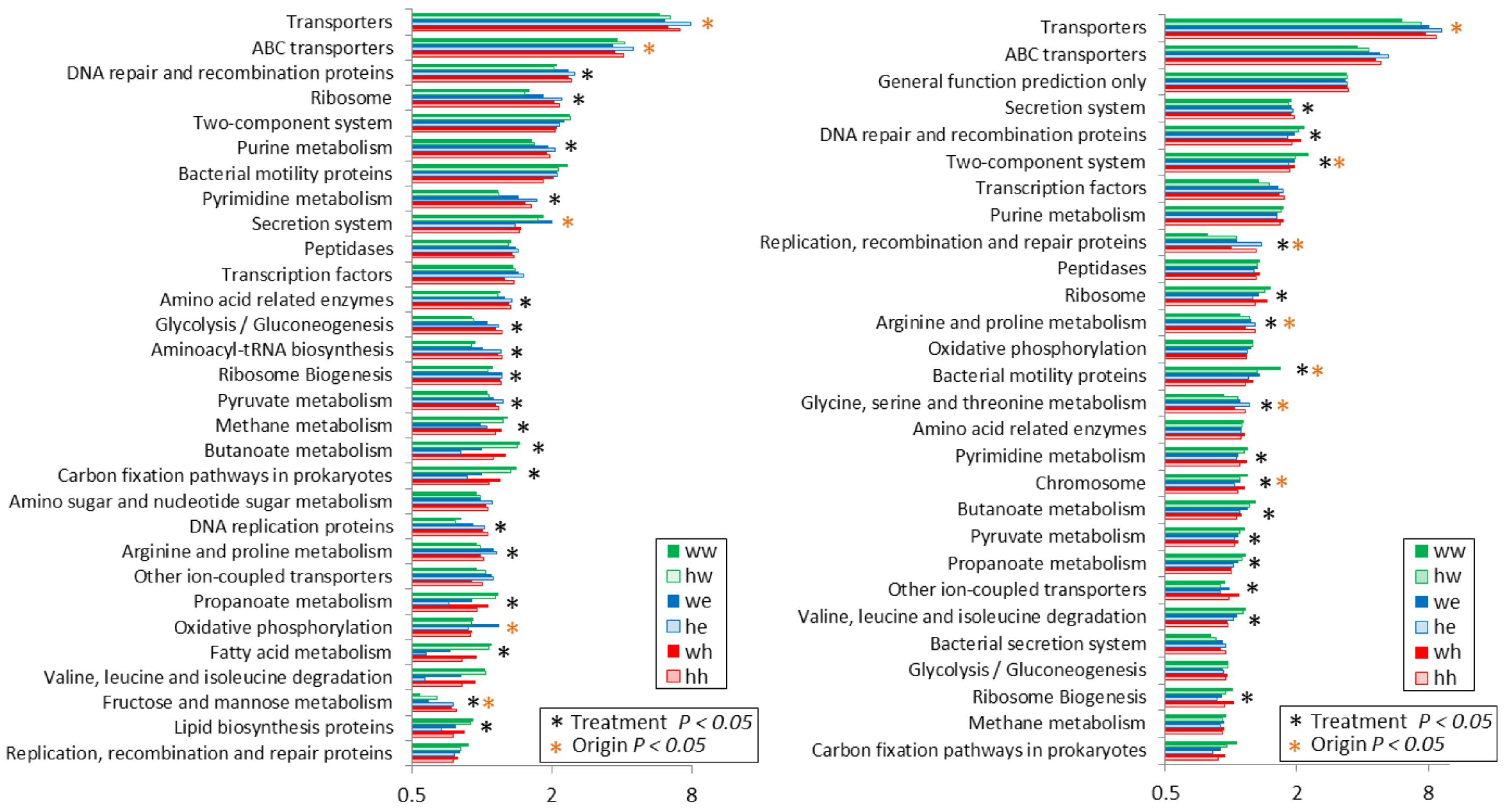


**Figure S5.** Functional analysis post-translocation, including the 30 most enriched KEGG pathways in the faeces and skin microbiomes. The terms differentially represented amongst environmental treatment groups, and between fish from different origins, are highlighted.
